# A novel method for producing functionalized vesicles that efficiently deliver oligonucleotides *in vitro* in cancer cells and *in vivo in mice*

**DOI:** 10.1101/2021.10.03.462960

**Authors:** Pragati Jain, Arthur G. Roberts

## Abstract

Nano-based delivery systems have enhanced our ability to administer and target drugs and macromolecules to their targets. Oligonucleotide drugs have great therapeutic potential but often have off-target effects and stability issues. Therefore, they are often encapsulated in vesicles with targeting ligands such as antibodies (Ab) to deliver their cargo. Herein, we describe a novel, scalable and straightforward approach to producing functionalized vesicles called the “Functionalized Lipid Insertion Method,” which differs from the older “Detergent-Dialysis Method.” The latter method required excess detergent and extensive dialysis over many hours to produce the functionalized vesicles. With our method, only the functionalized lipid is detergent-solubilized during the engineering of the vesicle. The approach reduces the dialysis time, keeps the vesicles intact while orienting the targeting moieties of the functionalized lipid toward the outside of the vesicle. Pilot *in vitro* and *in vivo* experiments was performed to show the feasibility of our method. Dynamic light scattering (DLS) experiments suggested that the original vesicular structure was relatively unperturbed, and the functionalized lipid was inserted externally. Our approach efficiently delivered oligonucleotides and affected the function of liver cancer HepG2 cells. Furthermore, functionalized vesicles achieved targeted delivery of oligonucleotides in mice without inducing a significant innate immune response. The industrial and therapeutic significance and implications of functionalized vesicles produced by our method are also discussed. Additional experiments and analyses are recommended to bring out the full potential of this molecular delivery technology.

## Introduction

Nano-based delivery systems have been developed to deliver a wide range of molecules, including drugs, nucleotides, and proteins [1]. Several nano-based delivery systems are available, including liposomes, dendrimers, and carbon nanotubes [2]. To be effective *in vivo* carriers, they are designed to be biodegradable, biocompatible, and non-immunogenic [3–5]. Ideal nano-based delivery systems must overcome many challenges, including rapid clearance, instability, toxicity, and inefficient targeting [5–7].

Two major nano-based delivery systems recently employed are extracellular vesicles (EVs) and liposomal nanoparticles (LNPs) [e.g., 8,9]. EVs are naturally occurring vesicles with an average diameter around 200 nm excreted from various body fluids such as blood and urine [10]. Depending on their source, their sizes range from 20 nm to 10 μm in diameter [10]. In contrast, LNPs are artificially produced and can be made into specific sizes through different techniques [e.g., 9,11]. EVs are formed by inward budding of the plasma membrane with other surface membrane invaginations from the Golgi apparatus [11]. These play an essential role in cell-to-cell communication and naturally carry RNA and proteins [11]. EVs are potentially advantageous for drug delivery because of their inherent biocompatibility, long-circulating half-life, low toxicity, and tendency to be endocytosed into target tissues [12]. On the other hand, LNPs also have a lot of therapeutic potential because they can be designed with improved biocompatibility, stored lyophilized for relatively long periods, and produced on an industrial scale [13,14].

EVs and LNPs can be designed to have a particular function by engineering them with a fluorescent fatty acid (FA) for tracking or a FA with a targeting molecule such as an antibody (Ab) or a receptor ligand. They can also be loaded with a range of molecules, from small molecules to proteins to oligonucleotides. These modified EVs (mEVs) and modified LNPs (mLNPs) fit into a general category of vesicles referred to as functionalized vesicles.

Functionalized vesicles have been engineered using relatively simple methods [15,16]. EVs have been covered with antibodies against the exosomal transmembrane protein CD9 and have improved delivery of miRNA to effector T-cells [15]. Since the early 1980s, an approach called the “Detergent-Dialysis Method” has been employed that uses detergent and dialysis to produce functionalized vesicles [17–24]. First, the process completely dissolves the vesicular components with detergent [17–24]. Then, these functionalized vesicles reassemble randomly by extensive and time-consuming dialysis to remove detergent [17–24]. Recently, a two-stage process for producing mEVs was developed where the EVs were PEGylated and covalently linked with antibodies [16]. The three-day process has a lot of potential for industrial scale-up [16,25,26]. Unfortunately, the mEVs undergo large temperature fluctuations between 4°C and 40°C, potentially destabilizing the cellular proteins of the mEVs and the attached antibodies (Abs) of the vesicles [16,25,26].

Functionalized vesicles can be loaded with oligonucleotide drugs. Oligonucleotide drugs can be from natural sources or manufactured synthetically [27,28]. They have shown a lot of potential for treating diseases, and several have been approved by the FDA [28]. Oligonucleotide drugs can be easily modified and produced, which is a major advantage over traditional small molecule drugs [29]. Most oligonucleotide drugs have sequences that complement their DNA or RNA targets [28]. Oligonucleotide drugs include small interfering RNAs (siRNA) [30,31], antisense oligonucleotides (ASOs) [32,33] and microRNAs (miRNAs) [34]. Unfortunately, oligonucleotide drugs suffer from immunogenicity, enzymatic degradation, and off-target effects [27].

SiRNA is generally recognized as a short 21 to 23 base pair double-stranded RNA (dsRNA) molecule with two 3’ nucleotide overhangs [30]. They play a vital role in RNA interference (RNAi), which is involved in cellular growth and differentiation [35]. They can also target several disease-causing genes [30,36]. Therefore, siRNA drugs represent viable drug candidates and are in different stages of drug development [30,36]. SiRNAs have been chemically modified and delivered through nanoparticle delivery vehicles such as liposomes [31].

ASOs are short 15-25 base pair polynucleotides that can be designed to bind to disease-related mRNA targets [32,33]. Mechanistically, they can attach to a target mRNA molecule and induce mRNA degradation [33]. Chemically modified ASOs can inhibit mRNA degradation by causing steric hindrance with their mRNA targets [33]. They ultimately affect mRNA translation and subsequent protein expression of the target mRNA molecule [33]. They have been delivered by various means, including liposomes, cationic amphiphiles, and dendrimers [33,37].

Finally, a small polynucleotide called microRNA (miRNA) represents another potential therapeutic that can be delivered with functionalized vesicles [34]. These macromolecules are small non-coding polynucleotides that range in length between 17 and 25 nucleotides [38]. They play an essential role in physiology by modulating gene expression through binding to mRNA [38]. MiRNAs have been associated with many diseases, including cardiovascular disorders, cancer, and allergic responses [39]. They can also elicit anticancer drug resistance in cancerous tumors [40]. Thus, miRNAs have served as biomarkers for diseases, and they can modulate gene expression [34,39]. These miRNA-related therapeutics can be synthetic mimics of naturally occurring miRNAs or miRNA inhibitors called anti-miRs [34]. Both miRNA mimics and anti-miRs have been chemically modified to reduce degradation and improve therapeutic efficacy [34]. They have also been encapsulated in liposomes, dendrimers, and polymers [34]. Several miRNA therapeutics are undergoing different drug development stages to treat cancer and liver diseases [34]. Therefore, targeted delivery of miRNA represents an excellent opportunity to test the effectiveness of our approach to produce functionalized vesicles.

In these pilot studies, we present a straightforward, efficient, and non-disruptive approach for producing functionalized vesicles called the “Functionalized Lipid Insertion Method.” The functionalized vesicles can be made much more efficiently than [16] and the older method [17–24]. Our innovative approach maintains the temperature where vesicles, proteins, and Abs are stable [25,26,41]. Functionalized vesicles can be engineered with diverse targeting molecules, including receptor ligands, antibodies, and affibodies. These vesicles will give them the potential ability to target any cell, tissue, or organ. The *in vitro* targeting and functional effects of mEVs and mLNPs by our procedure are demonstrated with liver cancer HepG2 cells. In mice, mEVs and mLNPs were designed with different Abs to target specific organs. The potential commercial advantages of targeted therapeutic delivery using our approach are discussed. In the last section, additional experiments and analyses are suggested to unleash the full potential of the “Functionalized Lipid Insertion Method” to deliver and study oligonucleotides.

## Materials and Methods

### Materials

The conjugating fatty acid (FA) label, 1,2-distearoyl-*sn*-glycero-3-phosphoethanolamine-*N*-[maleimide(polyethylene glycol)-2000] (DSPE-PEG2000-maleimide), and fluorescent 1,2-distearoyl-*sn*-glycero-3-phosphoethanolamine-N-(7-nitro-2-1,3-benzoxadiazol-4-yl) (NBD-DSPE) were purchased from Avanti Polar lipids (Alabaster, AL). HEPES (4-(2-hydroxyethyl)-1-piperazineethanesulfonic acid), *n*-dodecyl-β-*D*-maltoside (DDM) detergent, and Histopaque 1077 Reagent were purchased from Sigma Aldrich (St. Louis, MO). For the construction of liposomal nanoparticles (LNPs), *Escherichia (E.) coli* polar lipid extract was ordered from Avanti Polar lipids (Alabaster, AL), and chloroform was acquired from Sigma Aldrich (St. Louis, MO). Asialoglycoprotein receptor 1 (ASGPR1)/HL-1 polyclonal antibody (ASGR1_PAB_) (ab49355) and recombinant ACE2 monoclonal antibody (ACE2_MAB_) (ab108252) were purchased from Abcam (Cambridge, MA). NPHS2 (podocin) polyclonal antibody (NPHS2_PAB_) (MBS3013144) was acquired from MyBioSource (San Diego, CA). GFP monoclonal antibody (GFP_MAB_) (MA5-15256) was obtained from Thermo Fisher (Waltham, MA).

MiRNA mimics (mmu-miR-298-5p (0.38 mg/ml), hsa-miR-26a-5p (0.28 mg/ml)), and TaqMan™ MicroRNA assay were ordered from Thermo Fisher Scientific (Waltham, MA). Six ml Becton, Dickinson, and Company (BD) (Franklin Lakes, NJ) hematological tubes spray-coated with 1.8 mg/ml of dipotassium ethylene diamine tetraacetic acid (EDTA), and the blood separation agent Histopaque^®^ 1077 Reagent, which is a solution of polysucrose and sodium diatrizoate (1.077 g/mL), was obtained from Sigma-Aldrich (St. Louis, MO). DharmaFECT™ 4 transfection reagent was bought from Horizon Discovery (Cambridge, UK). Bio-Plex Pro Mouse Cytokine 8-plex Assay kit (#M60000007A) (Bio-Rad, Hercules, CA).

### Cell isolation and culture

Human peripheral blood mononuclear cell (PBMC) donors were enrolled for blood collection in compliance with the World Medical Association’s Declaration of Helsinki and the Human Research Protection Program and Institutional Review Board (IRB) guidelines for human subject research at the University of Georgia. In addition, healthy volunteers signed consent forms to inform them about the study. The human blood protocol (University of Georgia IRB no STUDY00006632) and the consent form were reviewed and approved by the IRB of the University of Georgia.

The human liver cancer HepG2 cells were purchased from American type culture collection (ATCC, Maryland, MD). These were grown and maintained in Eagle’s minimum essential medium (EMEM) (Corning), which was supplemented with 10% fetal bovine serum (FBS) (Atlanta Biologicals, Flowery Branch, GA) and 5% penicillin/streptomycin (Thermo Fisher, Waltham, MA). These cells were incubated in a humidified atmosphere of 5% CO_2_ at 37°C in Thermo Fisher Scientific Napco Series 8000 WJ CO_2_ incubator (Waltham, MA).

### Extracellular vesicle (EV) isolation

Approximately 10 ml of human blood were put into EDTA-coated BD hematological tubes to declot them. The peripheral blood nuclear cells (PBMCs) were isolated as described with some modifications [42,43]. Five ml of EDTA-treated blood samples were layered onto an equal amount of Histopaque^®^ 1077 Reagent in a 15 ml conical tube. The tube was centrifuged at approximately 400 g (1478 rpm) for 30 minutes and 4°C in an Eppendorf 5810R centrifuge (Hamburg, Germany), which separated the blood into plasma peripheral blood mononuclear cells (PBMCs) and erythrocyte layers. The top plasma layer was removed and discarded. The turbid middle layer of PBMCs was removed and put into a clean 15 ml conical tube. The tube was centrifuged at approximately 450 g (1917 rpm) for 10 minutes at 4°C. The supernatant was carefully removed using a transfer pipette to avoid disrupting the PBMC pellet. The pellet was then washed with 5 ml of an isotonic phosphate-buffered saline (PBS) (137 mM NaCl, 2 mM KCl, 10 mM Na_2_HPO_4_, 1.8 mM KH_2_PO_4_) solution and centrifuged twice at 300 g (1278 rpm). This pellet was suspended in RPMI growth media without glutamine and phenol red (Corning, NY, USA) and added to a 1M HEPES buffer solution (pH 7.4) to a final concentration of 25 mM HEPES to provide additional buffering capacity for the media. Cells were then transferred to a sterile Cellstar T-75 culture flask with a red filter screw cap (Greiner Bio-One, Monroe, NC) containing RPMI 1640 (with glutamine and phenol red) supplemented with 10% w/v fetal bovine serum (FBS) with 100 U/ml penicillin and 100 μg/ml streptomycin. This solution was incubated overnight (∼12 hours) in a humidified atmosphere at 37°C and 5% CO_2_ in Thermo Fisher Scientific Napco Series 8000 WJ CO_2_ incubator (Waltham, MA). Afterward, the media was transferred with a sterilized transfer pipette into 2 ml microcentrifuge tubes. The microcentrifuge tubes were centrifuged at 10,000 g (∼14,000 rpm) for 5 min. on a tabletop centrifuge at room temperature. About 1 ml of supernatant from each microcentrifuge tube was transferred to a new 2 ml microcentrifuge tube. Four hundred microliters of precipitation buffer B from the Qiagen (Formerly, Exiqon) miRCURY Exosome Isolation Kit (Qiagen, Germantown, MD) was added to the supernatant in each tube. The microcentrifuge tubes were inverted and vortexed, and incubated for ∼12 hours (overnight) at 4°C. The longer incubation time improved the EVs yield from the PBMCs. The remaining steps follow the manufacturer’s instructions for the Qiagen miRCURY Exosome Isolation Kit (Qiagen, Germantown, MD). The EVs were resuspended in 300 µl of the resuspension buffer supplied by the kit, combined in one tube, and stored at -80°C until needed. The total protein concentration of the EVs was measured using the Pierce Bicinchoninic Acid (BCA) Protein Assay Kit (Thermo Fisher Scientific, Waltham, MA) to estimate the concentration and the yield, which was typically around 10 μg.

### Preparation of liposomal nanoparticles (LNPs)

Unilamellar liposomal nanoparticles (LNPs) were prepared using the filter extrusion method [44]. The LNPs were composed of 80% w/v *E. coli* Avanti polar lipids and 20% w/v cholesterol as described previously [45]. Briefly, lipids and cholesterol were mixed in 10 ml of chloroform to get a final concentration of 10 mg/ml. This solution was evaporated to dryness in a Rotavapor Model R-114 (Buchi). After evaporation, the film was reconstituted in 0.1 mM EGTA and 50 mM Tris/HCl. This suspension was freeze-thawed at least ten times using liquid nitrogen and extruded 11 times through a LIPEX extruder (Northern Lipids, Burnaby, B.C., Canada) with a 400 nm cutoff Millipore filter (Millipore Sigma, Burlington, MA).

### Functionalized Lipid Insertion Method

A 200 μl fatty acid (FA) solution was made with 100 μM fluorescent NBD-DSPE and 100 μM DSPE-PEG2000-maleimide, and 0.1% w/v DDM detergent (10X the critical micelle concentration (CMC)) in an isotonic PBS (pH 7.4) buffer. The NBD-DSPE fluorescence was monitored at 550 nm by exciting at 445-460 nm using a SpectraMax M2 Plate Reader (Molecular Devices, Sunnyvale, CA). The FA solution was dialyzed using 0.5 mL Slide-A-Lyzer MINI Dialysis units with a 10 K_D_ cut-off filter (Thermo Fisher Scientific, Waltham, MA) against 2 L isotonic PBS for 2 h at 4°C to remove excess DDM detergent. Almost identical fluorescence for the NBD-DSPE FA was measured after dialysis showing that the lipids remained in solution after dialysis (data not shown). To ensure that all DSPE-PEG2000 maleimide were conjugated, two-fold excess of the targeting antibody (∼200 μM) was added to the dialyzed FA solution and incubated at room temperature for 1 hour. A 100 µl of purified 10 mg/ml EVs or LNPs was added to this 200 µl of the NBD-DSPE and DSPE-PEG2000-Antibody (DSPE-PEG2000-Ab) solution was briefly centrifuged to remove any large aggregates. The molar ratio of vesicle lipid to NBD-DPSE and DSPE-PEG2000-Ab was approximately 75:1:1. This 300 μl solution was incubated for 1 hour at room temperature to allow slow mixing and prevent any potential disruption of the vesicles. After incubation, the supernatant was dialyzed in a 0.5 mL 10 KD cut-off Slide-A-Lyzer MINI Dialysis unit against 2 L of isotonic PBS buffer for two hours at 4°C. Dialysis slowly removes the DDM detergent that surrounds the FAs. The dialysis process exposes hydrophobic surfaces of the derivatized or functionalized FA and entropically drives the FA tail into the vesicle bilayer to minimize exposure to water to form the functionalized vesicle. The inserted lipids, particularly the DSPE-PEG2000-Ab, remain embedded on the outer leaflet of the vesicle bilayer because a large thermodynamic barrier prevents them from flip-flopping to the inner leaflet of the bilayer [46]. A similar procedure has been used for directionally inserting membrane proteins into liposomal bilayers [52,53]. The orientation was confirmed enzymatically and through atomic force microscopy (AFM) [52,53]. Afterward, the mEV solution was incubated with 100 µl of precipitation buffer B from the Qiagen miRCURY Exosome Isolation Kit (Qiagen, Germantown, MD) overnight (∼12 hours) at 4 °C (Final volume = ∼250 μl). The mLNPs solution was then centrifuged at 14,000 rpm for 30 minutes to produce a pellet. To pellet the mEVs, the solution was centrifuged at 104,000 g (30,472 rpm) in a Beckman TLA 110 rotor for one hour at 20°C in a Beckman TLX ultracentrifuge. The supernatant was carefully removed. The functionalized vesicle pellet was then resuspended in 100 μl of isotonic PBS. The purpose of centrifuging the functionalized vesicles and removing the supernatant was to remove any remaining antibodies that were not cross-linked to the DPSE-PEG2000-Maleimide FA. The concentration of mEVs was measured by protein quantification using Pierce Bicinchoninic Acid (BCA) assay. The concentration of mLNPs was determined by tracking the amount of the lipid that was used throughout the procedure.

### Detergent-Dialysis Method

The production of modified vesicles by the “Detergent-Dialysis Method” was done as previously described [17–24]. All the components, the lipids, the proteins if present, and the derivatized lipids, were dissolved in detergent several times higher than the detergent’s CMC [17–24]. These solutions were then extensively dialyzed over many hours to remove the detergent [17–24]. The dialysis-driven detergent removal process causes the lipids, the proteins, and the derivatized lipids to randomly coalesce into modified vesicles of indeterminate sizes [17–24]. Differences between this method and the “Functionalized Lipid Insertion Method” are described in the manuscript text and Fig. 1.

**Figure 1.**
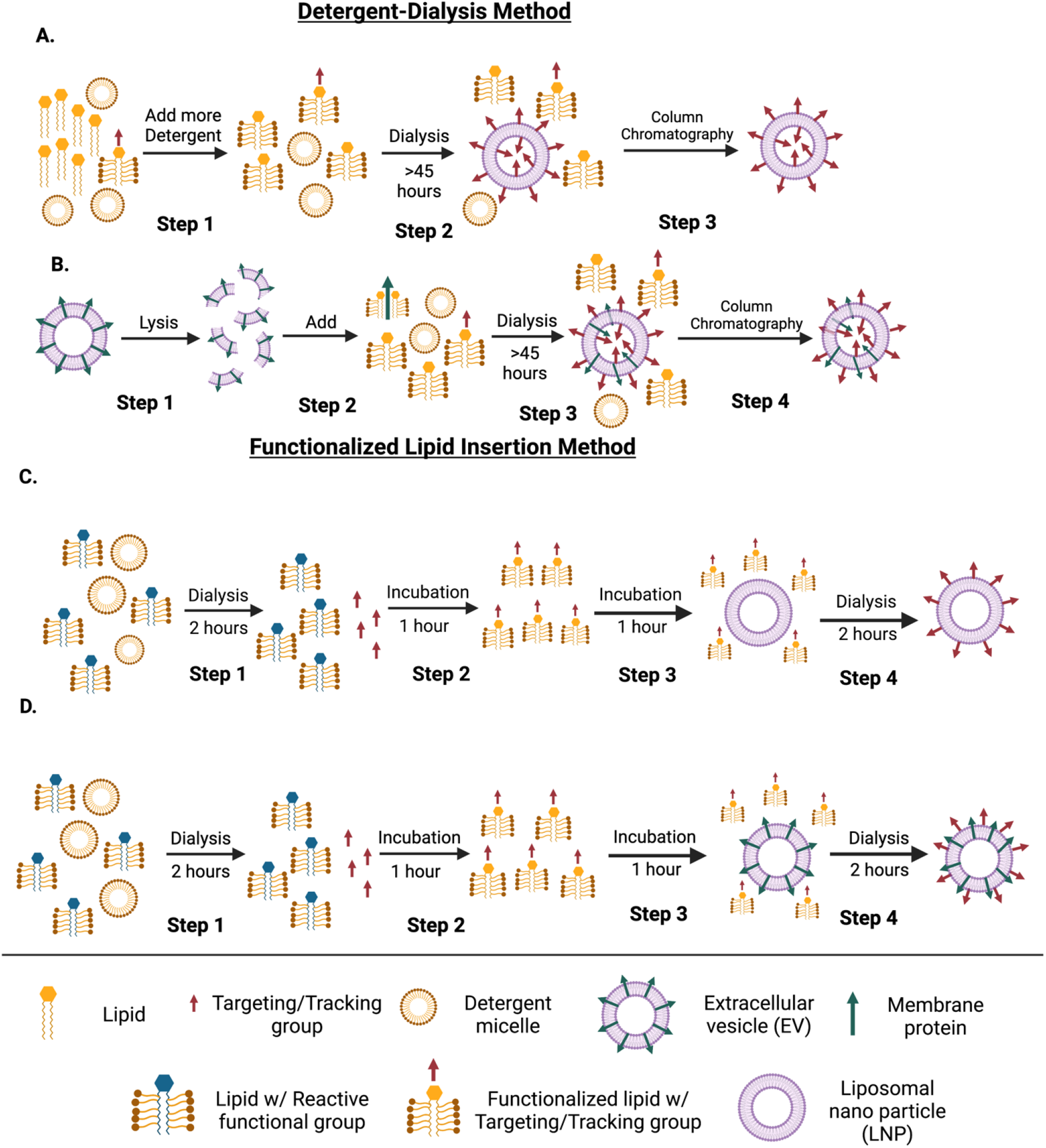
Bioengineering functionalized vesicles by the “Detergent-Dialysis Method” and the “Functionalized-Lipid Method.” A) Shows the “Detergent-Dialysis Method” that has been used since the early 1980s to produce functionalized vesicles from LNPs. In Step 1, functionalized lipids containing a targeting molecule such as an Ab are mixed with lipids and high concentrations of detergent, leading to complete solubilization of all the components. In Step 2, the high concentrations of detergent are dialyzed over time to remove the detergent. Once the detergent reaches a specific concentration, the mLNPs will spontaneously form, but the orientations of the functionalized lipids become randomized (red arrows) within the lipid bilayer. Even after this long dialysis period, column chromatography is often used to remove detergent-solubilized components in Step 3. B) Shows the same process of producing functionalized vesicles from natural vesicles like EVs. Because EVs are intact vesicles, they are lysed into their individual components in Step 1 prior to detergent solubilization and dialysis in Steps 2 and 3. In Step 3, because proteins are likely to be embedded in the natural vesicle lipid bilayer, their orientations will be randomly oriented (green arrows) like the functionalized lipids (red arrows). Column chromatography can be used in Step 4 to remove detergent-solubilized contaminants. C) Shows the process of producing mLNPs using the “Functionalized Lipid Insertion Method” that we developed. In Step 1, detergent-solubilized lipids with reactive functional groups are dialyzed for two hours to remove excess detergent. These solubilized lipids are mixed with targeting molecules such as antibodies (Abs) for 1 hour to form functionalized lipids in Step 2. In Step 3, the functionalized lipids are then incubated (or mixed gently) with preformed LNPs for another hour to ensure the integrity of the preformed vesicle. In Step 4, because the detergent concentration is relatively small compared with the detergent concentrations used in the “Detergent-Dialysis Method,” a lot shorter time is required for detergent removal. Removal of detergent from the functionalized lipids causes them to directionally insert into the preformed liposomes, becoming mLNPs. D) Shows natural vesicles like EVs being produced by the “Functionalized Lipid Insertion Method.” The process of producing mEVs is virtually identical to producing mLNPs. In Steps 1 and 2, a detergent-solubilized functionalized lipid is produced from dialysis and incubation with targeting molecules such as Abs. Then in Step 3, they are incubated (or gently mixed) with the EV to maintain vesicle integrity. Detergent removal through dialysis inserts the functionalized lipid directionally into the EV forming an mEV.

Lipids functionalized with antibodies (i.e., FA-PEG2000-Ab) were made as described above. About a milligram of the following mixtures was dissolved in 0.03% DDM by a similar procedure as in [17]: 1) *E. coli* Avanti polar lipids, 2) *E. coli* Avanti lipids, NBD-DSPE, and DSPE-PEG2000-Antibody with molar ratios of 75:1:1, 3) PBMC-derived EVs and 4) PBMC-derived EVs, NBD-DSPE, and DSPE-PEG2000-Ab with molar ratios of 75:1:1. These mixtures containing 0.03% DDM were put into 200 μl microcentrifuge tubes. Then they were sonicated in a Kendal ultrasonic cleaner HB23 (Kendal) at 25°C until the solutions clarified, indicating complete dissolution. The mixtures were then extensively dialyzed as described in [17] by putting 50 μl of them into 0.2 mL Slide-A-Lyzer MINI Dialysis units with a 10 KD cut-off filter (Thermo Fisher Scientific, Waltham, MA) against 2 L isotonic PBS for at least 45 hours at room temperature.

### Characterizing functionalized vesicles

Dynamic light scattering (DLS) was used to characterize the vesicle sizes by the “Detergent-Dialysis Method” and the “Functionalized Lipid Insertion Method” (Fig. 3), which is well established for this purpose [47–51]. Before the DLS experiments, all the samples were centrifuged at ∼8000 g (5,000 rpm) for 30 min. using a Microfuge™ 22R (Beckman Coulter, Brea, CA) at 4°C. The purpose of the centrifugation was to remove large particulates such as dust from the samples, which can interfere with the DLS measurement. Although not visible by the naked eye, all centrifuged samples were assumed to have a pellet, so only the top aqueous portion was carefully removed before DLS analysis. The DLS experiments were performed on a Malvern Zetasizer Nano ZS (Malvern Panalytical, Malvern, UK) using a Malvern 45 μl ultra-micro cuvette (ZEN2112). The DLS experiments were analyzed using the Zetasizer Software Version 8 (Malvern Panalytical, Worcestershire, United Kingdom), assuming a refractive index of 1.330 and a viscosity of 0.8872, which are parameters typically used for lipid-containing vesicles [52]. The size distribution curves in this manuscript were rendered on Igor Pro 6.3 (WaveMetrics, Portland, OR). Protein concentrations of the EVs were determined using the Pierce Bis-cinchonic assay kit (BCA), and LNPs were determined by tracking the lipid concentration throughout the experiments.

### Loading functionalized vesicles with miRNA

Electroporation was used to load miRNA in the vesicles, as it is an efficient method to load oligonucleotides into vesicles [48,53]. Equal amounts of mEVs or mLNPs were added with an equal amount of miRNA mimics (Thermo Fisher, Waltham, MA) in SFM to a total volume of 400 μl. The solution was put into 0.4 cm gap Bio-Rad (Hercules, CA) electroporation cuvettes. Functionalized vesicles were electroporated with 4 μg miRNA for the *in vitro* experiments and 100 μg miRNA for the *in vivo* experiments. Unmodified and modified vesicles were electroporated at 150 V and 10-15 ms with an exponential wave pulse in a Bio-Rad Gene Pulser X-Cell electroporator (Hercules, CA). Afterward, the samples were incubated at room temperature and at 4°C for 30 minutes to allow the vesicles to recover.

### miRNA delivery to HepG2 cells by functionalized vesicles

The human liver HepG2 cells were treated in various ways to determine their relative uptake of miRNA. The cells were counted using a hemocytometer on a Zeiss Invertoskop 40 C inverted microscope (Zeiss, Oberkochen, Germany). The cells were plated to ∼2 x 10^5^ cells per well in a 6-well VWR tissue culture-treated plates (VWR, Suwanee, GA) and incubated ∼12 hours (overnight). Afterward, the media was replenished with 2.7 mL of fresh 10% Eagle’s minimal essential medium (EMEM) media. The cells were FBS-starved for ∼12 hours before treatment to make proliferating HepG2 cells behave homogenously [54]. For experiments with unmodified and modified vesicles, 400 *µ*l of them were electroporated with 4 *µ*g miRNA and then they were added wells in the cell-containing 6-well plates. As per manufacturer’s instructions for the optimal siRNA transfection for the HepG2 cell line with 2 x 10^5^ cells/well, 8 μl of the DharmaFECT™ 4 transfection reagent was added with 4 *µ*g miRNA into a 400 μl solution. This solution was added to other cell-containing wells. The final volume for all the wells after adding the solutions was approximately 3 ml. Before RNA extraction, the treated plates were incubated for 72 hours under humidifying conditions at 37°C with 5% CO_2_ in a Thermo Fisher Scientific Napco Series 8000 WJ CO_2_ incubator (Thermo Fisher, Waltham, MA).

### RNA extraction and quantitative RT-PCR (qRT-PCR)

TRIzol Reagent (Invitrogen, Carlsbad, CA) was used to isolate intracellular RNA as described in the manufacturer’s protocol. All the isolated RNA was stored at -80°C. Complementary DNA (cDNA) was prepared from intracellular RNA (i.e., microRNA and the U6 small nuclear (snRNA)) using the TaqMan™ microRNA Reverse Transcription Kit (Thermo Fisher Scientific, Waltham, MA). The cDNA was analyzed on an Applied Biosystems 7900HT Fast Real-Time PCR System (Applied Biosystems, Foster City, CA) in a 384-well microplate with the TaqMan™ Universal PCR MasterMix (Thermo Fisher Scientific, Waltham, MA). For measurements to determine miRNA transfer efficiency, TaqMan™ MicroRNA assay (Thermo Fisher Scientific, Waltham, MA) kits with specific fluorescent cDNA primers for the miRNA and fluorescent cDNA primers for the U6 snRNA reference. The DNA was quantitated on the qRT-PCR instrument using the threshold cycle (C_T_) method with the U6 snRNA expression as a reference to calculate ΔΔC_T_ values, which correlates to the relative miRNA uptake [55–58]. These values were exported into Microsoft Excel format to calculate fold-difference and analyzed in GraphPad Prism 7 (GraphPad, San Diego, CA). The ΔΔC_T_ values were normalized against the non-coding U6 snRNA expression to indicate an approximate amount of miRNA delivered to the cells.

### *In-vitro* time course for miRNA uptake

Six-well VWR culture-treated plates (Suwanee, GA) were plated with HepG2 cells to a density of 100,000 cells/well. The HepG2 cells were counted using a hemocytometer on a Zeiss Invertoskop 40 C inverted microscope (Zeiss, Oberkochen, Germany). After the cells adhered, the cells were serum-starved for one day. The next day, the cells were treated with functionalized vesicles containing 4 μg of mmu-miR-298-5p. Before extracting RNA, the cells were incubated from 12 to 72 hrs in a humid atmosphere at 37°C with 5% CO_2_ in a Thermo Fisher Scientific Napco Series 8000 WJ CO2 incubator (Waltham, MA). Then, the RNA was extracted and analyzed by qRT-PCR using the procedures described above.

### Wound healing assay

For quantifying the functional effect of miRNA delivered by mEVs and mLNPs to HepG2 cells, the functionalized vesicles were loaded with a tumor suppressor miRNA hsa-miR-26a-5p [47,48]. The HepG2 cells were plated in a 24-well plate at a cell density of 3×10^5^ cells/well. A wound was created in each well using a 200 μl pipette tip followed by gentle washing with PBS buffer. The cells were then treated with mEVs and mLNPs containing miR-26a-5p at a vesicle dosage of 0.35 mg per well (i.e., 50 μl/well) in 500 μl of fresh 10% EMEM media and incubated for 72 hours. The controls for this experiment were untreated HepG2 cells (abbreviated Cells), cells treated with empty EVs (abbreviated Cells + EVs()), and cells treated with empty mEVs with the ASGR1_PAB_ (abbreviated mEVs(ASGR1_PAB_)) under similar conditions. The cells were imaged from 0 to 72 hours on an Olympus IX71 inverted Microscope with a TH4-100 power source (Tokyo, Japan). The images were analyzed using the Fiji image processing software, and the MRI wound healing tool in ImageJ Software (National Institutes of Health, Rockville, MD).

### Mouse Studies

Animal welfare at the University of Georgia is covered by the NIH Animal Welfare Assurance #: D16-00276/A3437-01. Our animal studies were performed under two animal use protocols (AUPs): A2018 03-025-R2 and A2020 03-014-A3. These studies were carried out at the animal facilities at the David Life Sciences Building and the Paul D. Coverdell Center at the University of Georgia in Athens, GA. We greatly appreciate the generosity of Drs. Mandi M. Murph and Yao Yao for providing some mice used in the studies and sharing their spaces for our animal studies. The strains, genders, ages, treatments of mice used for these studies are listed in Table S1 of the *Supplementary Information*.

### *In-vivo* Quantification of miRNA

After 72 hours, the mice were euthanized using carbon dioxide followed by a necropsy as described [59]. About 100 mg of tissue were obtained to assess the amount of miRNA delivered in the tissue. The sections were suspended in 1 ml of TRIzol reagent (Invitrogen, Carlsbad, CA, USA) in a 1.5 ml microcentrifuge tube. The tissues were homogenized with a 1000 µl digital pipette and a Bel-Art^TM^ Pro Culture Cordless Homogenizer Unit (Thermo Fisher, Waltham, MA). The samples were then centrifuged at 12,000 g for five minutes at 4°C to remove tissue debris with a Microfuge™ 22R centrifuge (Beckman Coulter, Brea, CA). The RNA was extracted using the Invitrogen PureLink™ RNA Mini Kit (Thermo Fisher, Waltham, MA) [60]. After RNA extraction, the RNA concentration was quantified using NanoDrop™ 2000/2000c Spectrophotometer (Thermo Fisher Scientific, Waltham, MA). The RNA concentration was determined using the extinction coefficient for single-stranded RNA in the spectrophotometer of 0.025 (μg/ml)^−1^ cm^−1^, and the RNA purity (>99%) was estimated by the 260 nm/280 nm ratio. The miRNA and U6 snRNA were amplified for quantification using the TaqMan™ microRNA reverse transcription and the TaqMan™ Universal PCR MasterMix (Thermo Fisher Scientific, Waltham, MA) as explained above.

### Cytokine assay to probe immunogenicity

The Bio-Plex Pro Mouse Cytokine 8-plex Assay kit was used to quantitatively determine the innate immune response of mice after 72 hours of functionalized vesicle treatment [61]. About a milliliter of blood was withdrawn immediately after euthanizing the animals in BD Microtainer^®^ tubes containing serum separator (SST^TM^) (Becton, Dickinson and Company, Franklin Lakes, NJ). The blood was allowed to clot at room temperature for 30 minutes. The tubes were then centrifuged at 1000 g for 15 minutes at 4°C using a Microfuge™ 22R (Beckman Coulter, Brea, CA), and the serum was then stored at -80°C. Before performing the assay, the serum was centrifuge at 10,000 g for 10 minutes at 4°C using a Microfuge™ 22R. The kit has immunoassays coupled to magnetic beads for detecting the following eight inflammatory factors, including the granulocyte-macrophage colony-stimulating factor (GM-CSF), interferon γ (IFN-g), tumor necrosis factor α (TNF-a), and several interleukins (IL), IL-2, IL-4, IL-5, IL-10, IL-12 (p70). The plate was then read using a Luminex MAGPIX system (Luminex, Austin, TX).

### Statistics

One-way analysis of variance (ANOVA) tests was used to determine statistical significance between groups comparing relative miRNA expression. A confidence interval of 95% with p-values significantly less than 0.05 being significant (*). Student’s T-Test was also used to compare two groups, with a 95% confidence interval. Data were analyzed with Microsoft Excel (Microsoft, Redmond, WA) and GraphPad Prism 7 (GraphPad, San Diego, CA).

## Results

### Methods for producing functionalized vesicles

This work describes a novel method for producing functionalized vesicles involving detergent and dialysis that we call the “Functionalized Lipid Insertion Method.” This method is distinct from the “Detergent-Dialysis Method” used for decades and described in the literature since the early 1980s [17–24].

The two approaches are shown schematically in Fig. 1. In the “Detergent-Dialysis Method” (Figs. 1A-B), all functionalized vesicle components are solubilized at a relatively high detergent concentration (Step 1) [e.g., 17,24]. The detergent-solubilized mixture must be extensively dialyzed over many hours or days (Step 2). As dialysis removes the detergent, the vesicles randomly form functionalized vesicles from the solubilized components. The composition of the original components will likely influence the functionalized vesicle size. During the process, the functionalized lipids in the mixture will randomly orient toward the inside and outside of the vesicle (red arrows near Step 3). However, the long dialysis period is often insufficient to remove all the unintegrated components and excess detergent. Therefore, column chromatography is often needed in addition to dialysis to remove contaminants that interfere with the functionalized vesicle (Step 3) [e.g., 17].

Fig. 1B shows the production of an mEV by the “Detergent-Dialysis Method.” A natural vesicle such as an EV must be broken up into its components before detergent-solubilization (Step 1) [19]. An undesirable side effect of this process is the complete disruption of the vesicular structure and possibly its natural functions. Next, the natural vesicle’s components, including lipids and proteins, and the functionalized lipid are dissolved at a high detergent concentration (Step 2). This detergent solubilization stage is followed by a long period of dialysis (Step 3). Because of very high detergent concentration, column chromatography is often needed to remove any remaining contaminating detergent or unintegrated components (Step 4). A vesicle composed of components from the original EV was reassembled randomly from the dialysis and column chromatography to form the mEV. As a result, both integral membrane proteins (green arrows) and functionalized lipids (red arrows) assume random orientations within the mEV. Some of the outward-facing proteins are now inward-facing proteins by this approach (green arrows). Some functionalized lipids face inside the vesicle cavity, where they cannot perform their intended function (red arrows).

A graphical representation of the “Functionalized Lipid Insertion Method” in this work is shown in Figs. 1C and 1D. The only component that is detergent-solubilized by this approach is the functionalized lipid. In the diagram, the reactive lipid (i.e., DPSE-PEG2000-Maliemide) is solubilized by detergent and dialyzed for two hours to remove excess detergent (Step 1). The purpose of eliminating the excess detergent is to prevent it from dissolving the target preformed vesicle. Then the detergent-solubilized reactive lipid is incubated for 1 hour with an excess of a targeting component like an antibody (Ab) to allow cross-linking between them (Step 2). Afterward, the functionalized lipid is incubated for another hour with a preformed artificial vesicle or a natural vesicle (Step 3). This incubation period allows the components to slowly combine without external perturbation that might disrupt the vesicles like sonication. Gentle mixing can be performed during this stage instead of incubation. Finally, the detergent bound to the functionalized lipid is removed by 2 hours of dialysis (Step 4). A lot less detergent is present, so the dialysis period is significantly shortened. Also, the relatively low detergent concentration reduces the risk that excess detergent will disrupt the target vesicle. Detergent removal from the functionalized lipid exposes hydrophobic parts of the FA tail. The functionalized lipid inserts itself into the target vesicle as an entropic process to reduce exposure to these hydrophobic surfaces. The FA tail of the functionalized lipid will insert into the target vesicle from the outside. Naturally, the functionalized part of the lipid on the other side of the molecule (red arrowhead) will be facing outward (Step 4). We have already experimentally demonstrated that we can orient integral membrane proteins in lipid bilayers using a similar principle [52,53].

### Possible mechanisms for functionalized vesicle endocytosis and miRNA delivery

The general theory for delivering molecules to a cell is that a targeting molecule on the functionalized vesicle will bind to a surface receptor or excreted receptor ligand. The vesicular uptake can occur by fusion or by several potential endocytosis mechanisms [62]. The specific mechanism of vesicular uptake has not been entirely resolved but appears to be cell-dependent [62].

Fig. 2 shows potential mechanisms that a functionalized vesicle can be endocytosed by a target cell. The top figure shows the binding of the PEG-linked Ab on the functionalized vesicle to a surface receptor on the cell (Step 1). The functionalized vesicle can bind to the receptor and induce endocytosis. Alternatively, the proximity of the functionalized vesicle to the plasma membrane may induce endocytosis through the assistance of the PEG linker (Step 2). Endocytosis of the receptor-bound vesicle leads to an invagination of the cell surface and eventually to the formation of an endosome (Step 3). The endosome eventually disintegrates intracellularly to release the miRNA or other cargo inside the target cell (Step 4).

**Figure 2.**
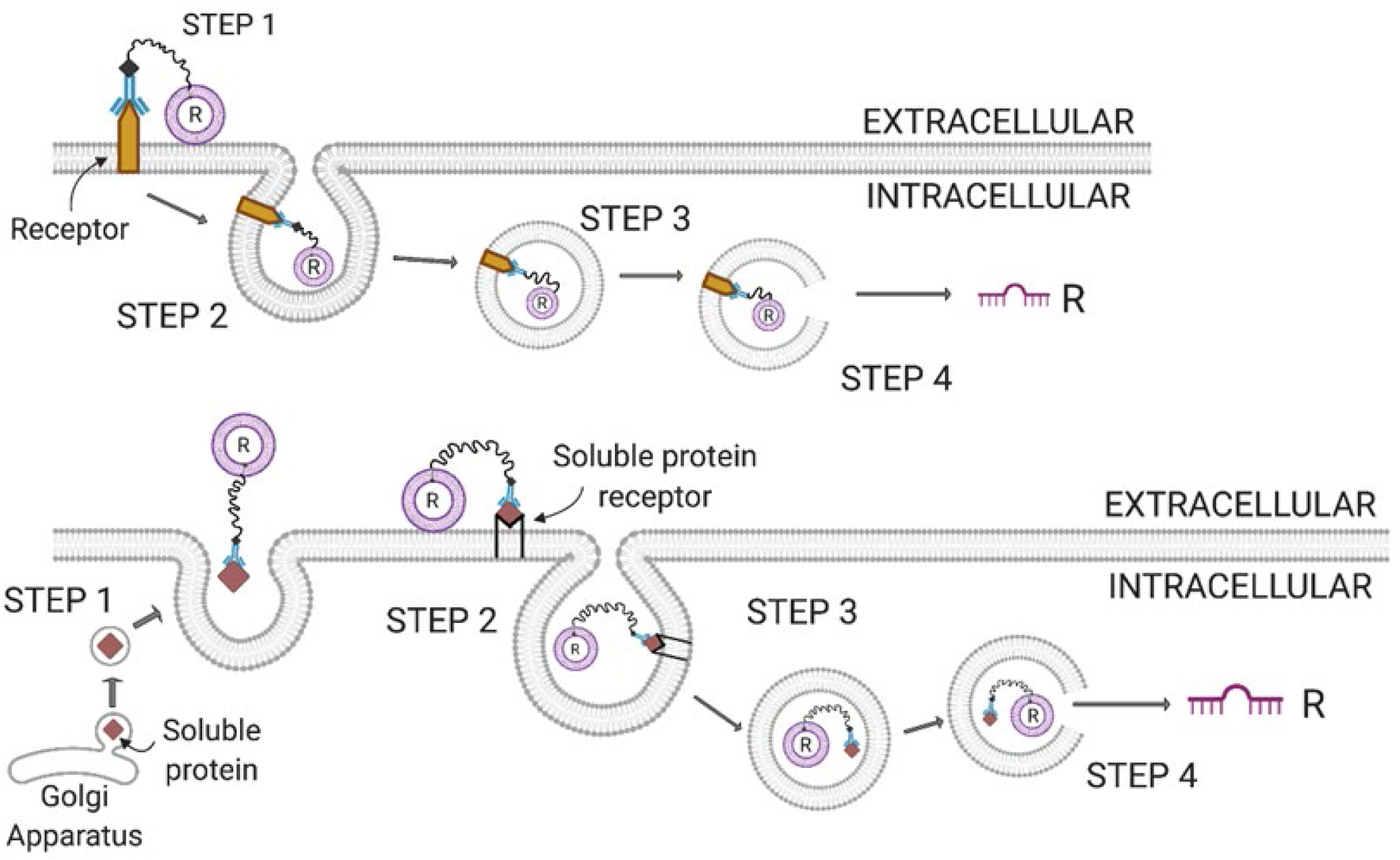
Potential mechanisms for cellular endocytosis of functionalized vesicles. A) A functionalized vesicle binds to a cell surface endocytotic receptor in Step 1 and then undergoes endocytosis in Step 2. The receptor-bound functionalized vesicle forms an endosome in Step 3 and eventually disintegrates in Step 4 to release its contents like the miRNA (R). B) A soluble receptor-ligand migrates to the plasma membrane surface from the Golgi apparatus, where it binds to a functionalized vesicle in Step 1. In order for the soluble protein-bound functionalized vesicle to be internalized by a cell, the complex binds to a cell surface receptor forming a ternary complex in step 2. If the surface receptor is endocytotic, the functionalized vesicle-bound receptor ligand is endocytosed to form an endosome in Step 3. Alternatively, if the functionalized vesicle is connected to the receptor-ligand through a long linker like PEG2000, the functionalized vesicle might contact the plasma surface leading to endocytosis. The resulting endosome disintegrates and releases its contents, such as R in Step 4. Abbreviations: R, miRNA.

The bottom of Fig. 2 shows the targeting of secretory receptor ligands from cells by functionalized vesicles containing miRNA. In Step 1, soluble proteins are trafficked to the Golgi apparatus from the endoplasmic reticulum (ER) [63]. The Golgi apparatus produces secretory vesicles containing the receptor ligands that migrate to the plasma membrane [63]. At the plasma membrane, they are secreted into the extracellular (EC) space, where they are bound by a functionalized vesicle (Step 1) [63]. The functionalized vesicle bound-receptor ligands bind to cell surface receptors on the plasma membrane forming a ternary complex with the receptor. Receptor-ligand binding to the surface receptor may lead to conformational changes that lead to endocytosis (Step 2) [e.g., 64]. The functionalized vesicle is tethered to the receptor-ligand and surface receptor by a PEG linker allowing the functionalized vesicle to interact with the plasma membrane surface. Alternatively, the interaction of the functionalized vesicle through the PEG linker with the plasma membrane can lead to endocytosis. In either case, the functionalized vesicle will be engulfed by the cell to form an endosome (Step 3) [62]. The endosome containing the functionalized vesicle loaded with miRNA migrates within the cell (Step 3). Eventually, the endosome and the vesicle disintegrate, releasing the miRNA or other cargo intracellularly (Step 4).

### The dynamic light scattering (DLS) technique reveals how different processes affect vesicle size

The DLS technique is a well-established technique for estimating particle sizes based on their dynamic light scattering properties [65]. The technique is well suited to estimate particle size distributions [e.g., 50,66]. The DLS technique was used to determine the effects of the “Detergent-Dialysis Method” and the “Functionalized-Lipid Insertion Method” on the vesicle size distributions. Because functionalized vesicles are assembled all at once with the “Detergent-Dialysis Method,” these experiments will test the hypothesis that the two approaches will influence the vesicle size distribution.

Fig. 3 shows DLS measurements of functionalized vesicles produced by the two methods. The predominant particle sizes were determined in each solution by comparing the particle size distribution (PSD) to the particle diameter. Fig. 3A shows the predominate DDM micelle size (solid line) and a detergent-solubilized NBD-DSPE or DPSE-PEG2000-maleimide micelle size (dotted line) present in solution. The DDM micelle particle size distribution (PSD) was 6.8 ± 1.6 nm in diameter. In comparison, the detergent-solubilized FA PSD was slightly smaller at 5.7 ± 1.4 nm, consistent with the diameters you would expect for these molecules.

**Figure 3.**
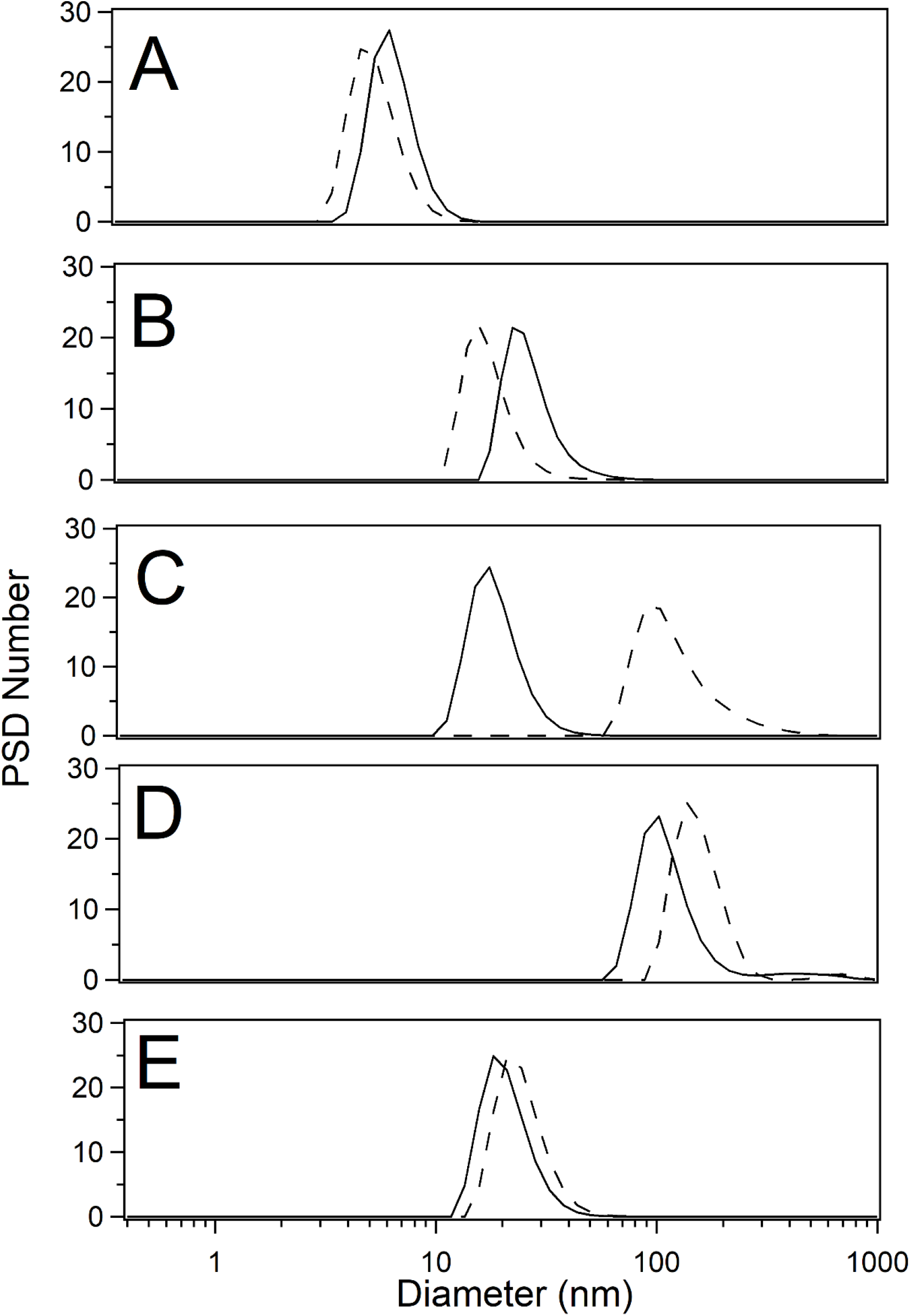
Representative Size distribution of functionalized vesicles determined by dynamic light scattering (DLS). A) Detergent DDM micelles (solid line) and DDM-solubilized FAs (dashed line). The detergent solubilized FAs are an equal mixture of NBD-DSPE and DSPE-PEG2000-maleimide. The “Detergent-Dialysis Method” was used B) to produce the size distributions of LNPs (solid line) and mLNPs (dashed line) and C) to produce the size distributions of EVs (solid line) and mEVs (dashed line). The solid lines in panels D) and E) show the size distributions produced for LNPs using the extrusion approach [44] and EVs through isolation and purification protocols described in the *Materials and Methods*. The dashed lines in panels D) and E) show the size distributions produced for mLNPs and mEVs using the “Functionalized Lipid Insertion Method” described in the text.

Figs. 3B and 3C show vesicles produced using the “Detergent-Dialysis Method.” In Fig. 3B, the LNP (solid line) had an average diameter of 87.8 ± 34.7 nm, similar to previous observations [20]. However, the mLNP (dotted line) had about half the diameter of the LNP at 51.2 ± 19.5 nm. This size difference suggests that the size of LNPs produced by this method may be sensitive to their initial component mixtures, as we hypothesized. Fig. 3C shows the EV (solid line) and mEV (dotted line) PSDs produced using the “Detergent-Dialysis Method.” The EVs created using this method had an average diameter of 19.7 ± 5.6 nm. In contrast, the average diameter of mEVs produced by this approach increased 6-fold larger or 131 ± 64.5 nm, showing again how the initial component mixtures can influence vesicle size.

Fig. 3D and Fig. 3E show the unmodified vesicles and modified vesicles produced using the “Functionalized-Lipid Insertion Method.” Fig. 3D shows the PSD of LNPs that were unmodified (solid line) and had an average diameter of 115.6 ± 35.0 nm. The average diameter of mLNPs (dotted line) increased ∼30% to 160.3 ± 37.4 nm. A 40 nm diameter increase is consistent with the length of the PEG2000 linker and Ab connected to the mLNP [67,68]. The unmodified EV had an average diameter in Fig. 3E (solid line) of 21.2 ± 5.9 nm, which is similar to the EV sizes produced with the “Detergent-Dialysis Method” (Fig. 3C, solid line). In Fig. 3E (dotted line), the average diameter of EVs increased only 17% or 5 nm to 24.7 ± 6.9 nm. The modest diameter increase for mEVs implied that their original structures were intact, but less functionalized lipid was inserted per EV than LNPs. One possibility for the size difference between mLNPs and mEVs is that the plasma membranes of EVs are crowded with many intrinsic membrane proteins that interfere with functionalized lipid insertion [e.g., 69]. The other possibility is geometrical. Because mEVs are smaller than mLNPs, their surface areas will be much larger within a given volume than mLNPs. In other words, a lot more functionalized lipids will be required to saturate the membrane surface of mEVs than mLNPs.

### *In-vitro* delivery of miRNA by functionalized vesicles produced from the “Functionalized-Lipid Insertion Method”

The human liver HepG2 cancer cell line has been the focus of many *in vitro* studies related to the functioning of liver cells and liver cancer [70–72]. The endocytotic asialoglycoprotein receptor 1 (ASGR1) is highly expressed on the surface of HepG2 cells [73]. The endocytotic process involves receptor-ligand binding and ASGR1 trimerization through receptor-mediated endocytosis [74]. This receptor has already been exploited to mediate small molecule uptake into HepG2 cells [75–77]. For example, galactosamine-linked albumin and nanoparticles loaded with pullulan and arabinogalactan delivered the anti-cancer drug doxorubicin to HepG2 cells [75,76]. In another study, PEGylated liposomes modified with lactoferrin successfully delivered the fluorophore coumarin-6 through ASGR1 receptor-mediated endocytosis of HepG2 cells and HepG2 cells implanted in nude mice [77]. Finally, ASOs have been delivered to HepG2 cells by linking them to the ASGR1 receptor substrate *N*-acetylgalactosamine [78]. As far as we know, ASGR1_PAB_ or any ASGR1 Abs have never been used to facilitate molecular uptake through the ASGR1 receptor with HepG2 cells. In this study, functionalized vesicles were made with ASGR1_PAB_, which targets the EC domain of the ASGR1 receptor.

The focus of my laboratory is on drug transporters called P-glycoprotein (P-gp), which plays a major role in drug disposition, anti-cancer drug resistance, and biological barriers like the blood-brain barrier [79]. Despite being studied for decades, transporter measurements with P-gp and other transporters remain challenging [80]. The P-gp expression can vary between different cancer cell lines used to measure transport [81]. Even variants of the same mammalian cancer cell line can have significantly different properties [82]. Furthermore, transporter transfection of mammalian cells can affect the expression levels of other transporters within the transfected cell [83], further complicating transport analysis. In addition, control mammalian cancer cells without P-gp can express different endogenous transporter levels than the mammalian cancer cells transfected with P-gp [81]. Finally, even culturing conditions can significantly affect P-gp expression levels [84]. Therefore, a critical need exists to keep drug transporter expression levels constant.

We were hopeful that oligonucleotides targeting P-gp and delivered by functionalized vesicles could control drug transporter levels. Our former collaborator, a miRNA expert, had us investigate P-gp and functionalized vesicles with mouse miR-298 based on [85,86], which they indicated affected P-gp expression. The miRNA affecting P-gp expression was essential to use our NIH funding, which is focused on P-gp and funded this work. Unfortunately, the miR-298 in [85,86] was human miRNA-298, with completely different targeting properties and biological functions than mouse miRNA-298 [87]. As far as we know, mouse miRNA-298 is not known to affect P-gp expression in any non-primate species significantly. That revelation was one of the primary reasons that the collaboration was terminated. Despite this setback, we felt the described functionalized vesicle technology could eventually be used to investigate P-gp. We also felt obliged to provide the procedure and data to the scientific community with minimal additional NIH funds to demonstrate feasibility. The mouse miRNA-298 was used to test the feasibility of functionalized vesicles since it would theoretically not have any physiological effects on the human liver HepG2 cells.

MiRNA nomenclature can be confusing for a novice. The formal names for mouse and human miR-298 are mmu-miR-298-5p and hsa-miR-298-5p, respectively [87]. Because of the similarity of their names, we were led to believe that they had similar functions by our former collaborator. The first three letters are the species [87]. Mmu is for *Mus (M.) musculus* (house mouse) and hsa is for *Homo* (*H.*) *sapiens* [87]. The number typically reflects the order of miRNA deposition into the miRNA database called miRbase and is almost always unrelated to the biological function [87]. The 5p refers to the 5’ side of the precursor miR-298 stem-loop that the miRNA comes from [85]. The mouse miR-298 step loop has a second miRNA on the 3’ side called mmu-miR-298-3p, which can be functionally distinct from the miRNA on the 5’ side [88].

Fig. 4 are experiments to estimate the relative uptake of mmu-miR-298-5p in HepG2 cells. The experiments shown in Fig. 4A were used to gauge the uptake of 4 μg of mmu-miR-298-5p into the HepG2 cell line under different conditions. The relative uptake of mmu-miR-298-5p was normalized against HepG2 cells treated with DharmaFect 4 transfection reagent with miRNA (abbreviated DharmaFect(miRNA)). No mmu-miR-298-5p was detected in untreated HepG2 cells. HepG2 cells treated with EVs electroporated with mmu-miR-298-5p (abbreviated EVs(miRNA)) had miRNA uptake efficiency that was 5-fold higher than HepG2 cells treated with DharmaFect(miRNA). The miRNA uptake efficiency of HepG2 with EVs constructed with the ASGR1_PAB_ and loaded with mmu-miR-298-5p (abbreviated mEVs(ASGR1_PAB_, miRNA) showed 8-fold greater improvement in miRNA uptake than the DharmaFect(miRNA) control. In contrast, HepG2 cells treated with exosomes genetically engineered with an Apolipoprotein A1 targeting ligand had more efficient miRNA uptake [48]. The Apolipoprotein A1 targeting ligand on the exosome may function better with the ASGR1 receptor than the ASGR1_PAB_ because they target different locations on the receptor’s extracellular domain (EC) domain [74]. Also, Apolipoprotein A1 naturally binds in a manner that initiates receptor-mediated endocytosis [74,89,90]. Modified LNPs with ASGR1_PAB_ were also made and loaded with 4 μg mmu-miR-298 (abbreviated mLNP(ASGR1_PAB_, miRNA)). Relative miRNA uptake was almost 2-fold higher in HepG2 cells treated with mLNPs (ASGR1_PAB_, miRNA) than cells treated with mEVs (ASGR1_PAB_, miRNA). The higher miRNA uptake by HepG2 cells treated by mLNPs may be due to more functionalized lipids per LNP or more miRNA delivered per vesicle because of their relatively large size compared to EVs.

**Figure 4.**
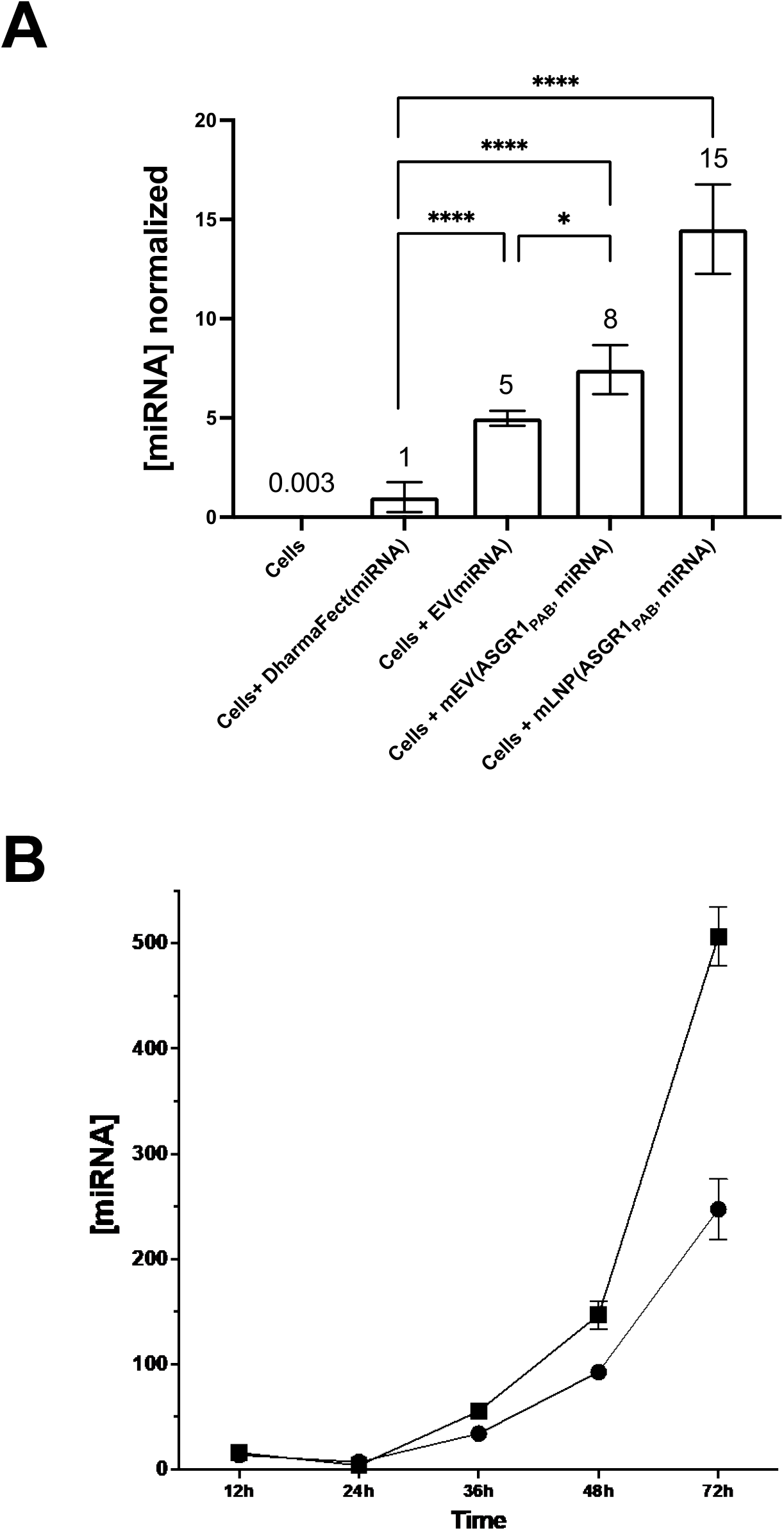
Relative miRNA uptake efficiency into HepG2 cells with different treatments. A) The relative mmu-miR-298-5p (miRNA) uptake into HepG2 cells under the various treatments: Untreated HepG2 cells (Cells), HepG2 cells treated with DharmaFect 4 transfection reagent and miRNA (Cells + DharmaFect(miRNA)), HepG2 cells treated with EVs and miRNA (Cells + EVs( miRNA)), HepG2 cells treated with mEVs with an ASGR1_PAB_ and containing miRNA (Cells + mEVs(ASGR1_PAB_, miRNA), and HepG2 cells treated with mLNPs with an ASGR1_PAB_ and containing miRNA (Cells + mLNPs(ASGR1_PAB_, miRNA). The relative miRNA uptake was normalized against the miRNA uptake by treatment with DharmaFect(miRNA). The values in the panel represent the mean ± SD of at least three independent experiments. B) Temporal changes in miRNA uptake efficiency for mLNPs(ASGR1_PAB_, miRNA) (solid squares) and mEVs(ASGR1_PAB_, miRNA) (solid circles) over 72 hours with respect to U6 snRNA. The points and bars represent the average and SD, respectively. The p-values are * p ≤ 0.05, ** p ≤ 0.01, *** p ≤ 0.001, **** p<0.0001. Abbreviations: SD, standard deviation.

The miRNA uptake’s temporal efficiency depends on the miRNA endocytotic mechanism, the miRNA carrier vehicle, and the miRNA type [91,92]. For example, the miRNA uptake of a human ovarian cancer cell line treated with liposomes loaded with fluorescently-labeled hsa-miR-7-5p increased linearly over 24 hours [91]. The uptake of miRNA in multiple myeloma treated with stable nucleic acid-lipid particles (SNALPs) loaded with hsa-miR-34a decreased after 48 hours [92]. In these pilot experiments, the miRNA uptake in HepG2 cells after treatment with mLNP(ASGR1_PAB_, miRNA) and mEV(ASGR1_PAB_, mRNA) was examined over 72 hours (Fig. 4B). The uptake efficiency of HepG2 cells to mmu-miR-298-5p increased exponentially by treatment with mLNPs or mEVs. After incubating the cells for 24 hours, the uptake efficiency doubled every ∼12 hours for the mEVs and ∼8 hours for the mLNPs. The relative miRNA uptake at 72 hours for mLNPs and mEVs was 600 and 250-fold higher, respectively, above untreated HepG2 cells versus the U6 snRNA. The relatively high miRNA uptake at 72 hours was deemed a detectable miRNA uptake and an appropriate incubation time to test mEVs and mLNPs in mice.

### Modified vesicles reduce HepG2 cell proliferation by enhancing hsa-miR-26a-5p uptake

The functional effects of treating HepG2 cells were tested by treatment with the mEVs(ASGR1_PAB_) and mLNPs(ASGR1_PAB_) loaded with miRNA hsa-miR-26a-5p (abbreviated mEV(ASGR1_PAB_, miRNA) and mLNP(ASGR1_PAB_, miRNA)). The miRNA hsa-miR-26a-5p was chosen because it inhibits HepG2 cell proliferation and promotes HepG2 cell apoptosis [48]. In other words, the miRNA will slow the closure gap of wounds made from a layer of HepG2 cells.

Wound healing assays with bright field microscopy has been effectively used to gauge cell proliferation and migration [93,94]. The approach was used to estimate the effect of mEVs and mLNPs treatment of HepG2 cells in Fig. 5. Fig. 5A shows the wound closures of HepG2 cells after various treatments at 0 and 72 hours. The wound closed to about half its original width for untreated cells, indicating uninhibited HepG2 cell proliferation. The HepG2 cells treated with empty EVs (abbreviated EV()), unmodified extracellular vesicles electroporated with hsa-miR-26a (abbreviated EV(hsa-miR-26a)), and empty modified extracellular vesicles with the polyclonal antibody for the ASGR1 receptor (abbreviated mEV(ASGR1_PAB_)) had an almost similar extent of wound closure as the untreated cells meaning no effect on HepG2 cell proliferation. The wound area for HepG2 cells after treatment with mEVs(ASGR1_PAB_, miRNA) and mLNPs(ASGR1_PAB_, miRNA) almost completely inhibited wound closure, demonstrating efficient uptake of hsa-miR-26a.

**Figure 5.**
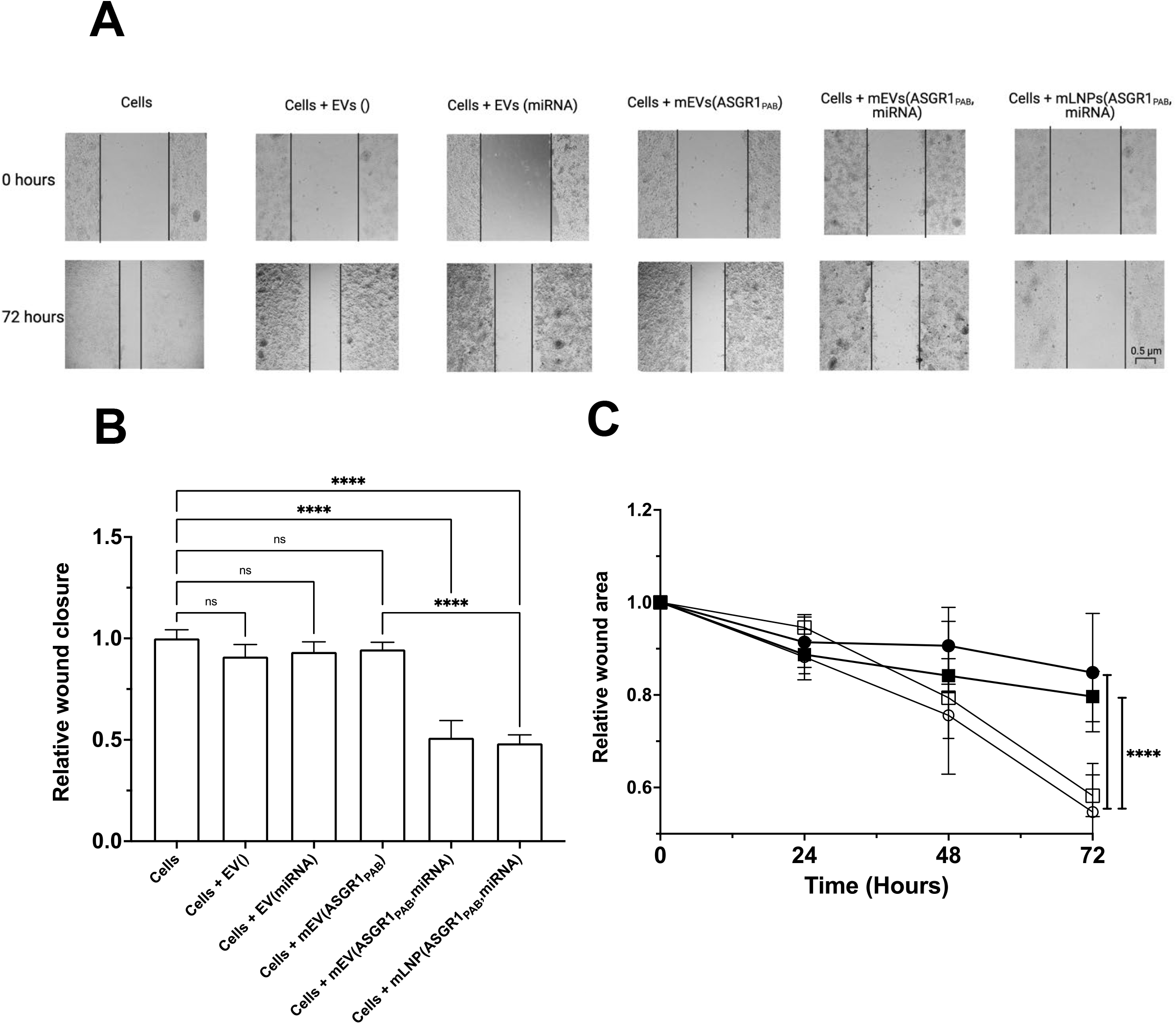
Anti-proliferative effects miRNA hsa-miR-26a-5p with HepG2 cells under various treatments. A) Microscopic images of a HepG2 layer during the migration assay with the following treatments after 0 (top) and 72 hours (bottom): Untreated HepG2 cells (Cells), HepG2 cells treated with empty EVs (Cells + EVs()), HepG2 cells treated with EVs containing miRNA (EVs(miRNA)), HepG2 cells treated with empty mEVs engineered with an ASGR1_PAB_ (Cells + mEVs(ASGR1_PAB_)), HepG2 cells treated with mEVs with the ASGR1_PAB_ and miRNA (Cells + mEVs(ASGR1_PAB_, miRNA)), and HepG2 cells treated mLNPs with the ASGR1_PAB_ and miRNA (Cells + mLNPs(ASGR1_PAB_, miRNA)). The black square bracket in the lower right-hand image represents a distance of 0.5 μm. B) Shows quantitative measurements of HepG2 cell migration using the data and same abbreviations from panel A. The values in the bar graphs represent the mean ± SD of at least three independent experiments. C) Temporal changes in HepG2 cell migration in untreated cells (open circles) and cells treated with EVs(miRNA) (open squares), mEVs(ASGR1_PAB_, miRNA) (closed circles), and mLNPs(ASGR1_PAB_, miRNA) (closed squares) over 72 hours. To differentiate the functionalized vesicles, the time course for HepG2 cells treated with mEVs(ASGR1_PAB_, miRNA) and mLNPs(ASGR1_PAB_, miRNA) is shown as thick black lines. The error bars represent the mean of three independent experiments ± SD. The dose miRNA in each of the experiments was 0.35 μg. The p-values are * p ≤ 0.05, ** p ≤ 0.01, *** p ≤ 0.001, **** p<0.0001. Abbreviations: SD, standard deviation.

This wound closure over 72 hours was assessed quantitatively in Fig. 5B. The percent wound closure by all the treatment groups was normalized to the amount of wound healing in the untreated cells. The results show that the relative amount of wound closure in the cells treated with empty EVs(), EVs(hsa-miR-26a), and mEVs(ASGR1_PAB_) was virtually identical to untreated cells. For EVs(hsa-miR-26a), the lack of effect on cell proliferation may be due to inefficient uptake of the hsa-miR-26a miRNA by the HepG2 cells. In contrast, wound closure by cells treated with mEVs(ASGR1_PAB_, hsa-miR-26a) or mLNPs(ASGR1_PAB_, hsa-miR-26a) inhibited cell proliferation by 50%.

Along with wound closure, the relative wound area was also analyzed temporally in Fig. 5C. The wound in untreated HepG2 cells closed at 16 ± 3 % per day (open circles). This wound closure rate is close to the 19% per day observed for HepG2 treated with extracellular vesicles containing hsa-miR-26a (calculated from Fig. 5B in [48]). After three days, the wound area was reduced to 52 ± 8% of the original area, suggesting uninhibited cell proliferation. The relative wound area in the cells treated with EVs delivering miRNA (abbreviated EVs(miRNA), open squares) was reduced at the rate of 13.9 ± 1.5 % per day, thus not differing significantly from untreated cells. Treating the HepG2 cells with mEVs(ASGR1_PAB_, miRNA) decreased the rate of closure to 5 ± 4.3% per day (closed circles) and with mLNPs(ASGR1_PAB_, miRNA) reduced the rate of closure to 6.7 ± 1.8% per day (closed squares). HepG2 cells treated with either our mEVs or mLNPs inhibited cell proliferation better than HepG2 cell treatment with miRNA-loaded Apo A1 targeting ligand exosomes, which only decreased HepG2 cell proliferation to 10% per day (calculated from Fig. 5B in [48]).

### Process of *in vivo* targeting of modified vesicles

The uptake of miRNA in mice after treatment with mEVs and mLNPs is shown in Fig. 6. Modified vesicles are loaded with miRNA by electroporation and injected intraperitoneally (IP) into mice (Fig. 6A). Because of their small physical size, IP administration of mice was easier to perform and more reliable than intravenous (IV) administration [95–99]. Also, vesicles administered by IP injection distribute broadly to the organs and the visceral adipose tissue because of access to the lymphatic system [95–99].

**Figure 6.**
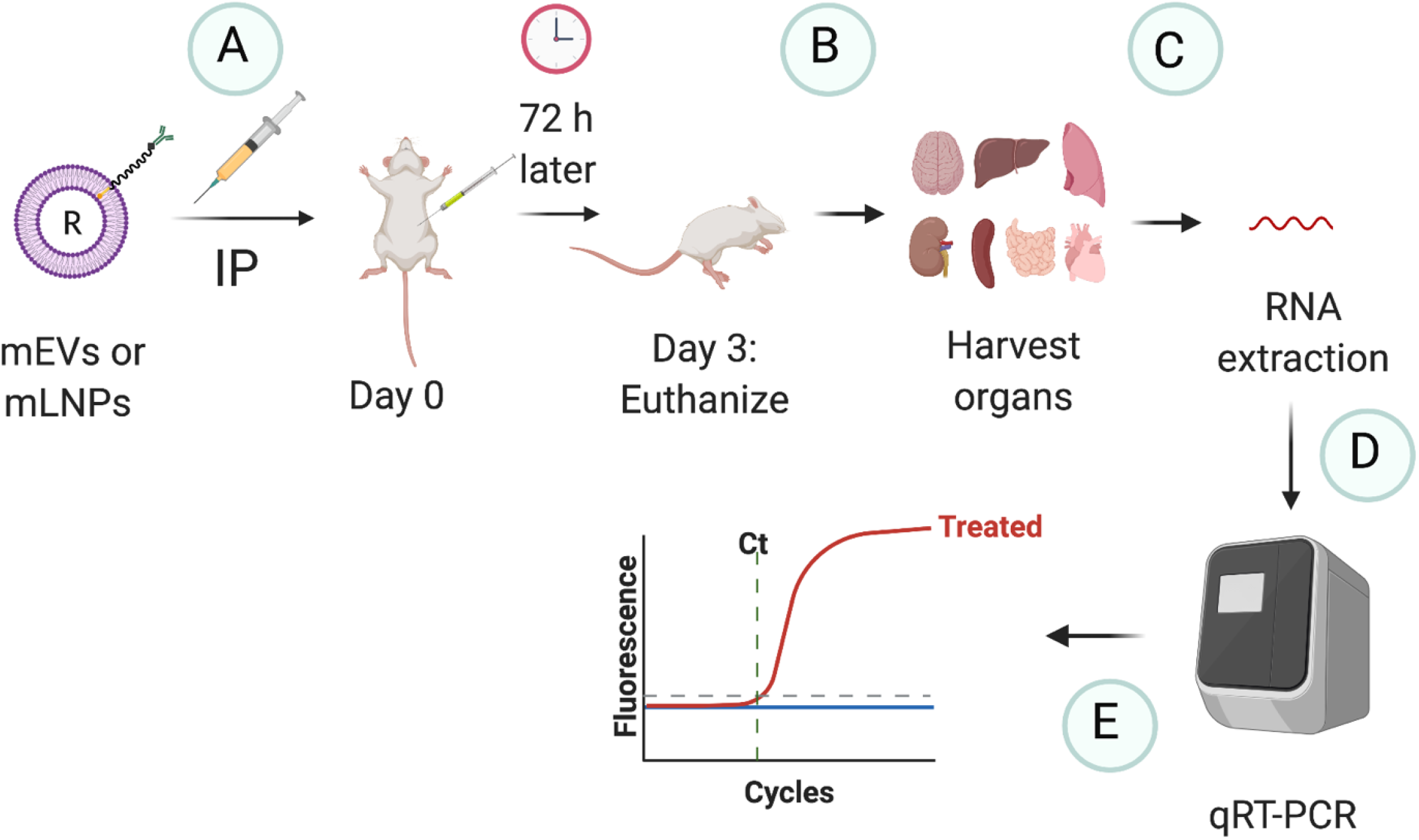
Process of administering functionalized vesicles into mice. A) On the first day, functionalized vesicles, mEVs or mLNPs, with miRNA (R) and targeting ligands such as Abs are intraperitoneally (IP) injected into mice. B) Three days later, the mice are euthanized, and the organs are harvested. C) RNA is harvested and purified from the mouse organs. D) The amount of purified RNA from each of the organs is analyzed against U6 snRNA by qRT-PCR. E) The relative level of miRNA uptake by the mouse organs from treatment with the functionalized vesicles is determined using the ΔΔC_t_ method. Abbreviations: R, miRNA.

The treated mice were euthanized after three days using carbon dioxide followed by cervical dislocation, and their organs were harvested (Fig. 6B). First, the tissue was analyzed using RNA extraction protocols described in the *Materials and Methods* and Fig. 6C. Then the sigmoidal amplification of miRNA and U6 snRNA in the mouse organs was measured in real-time by fluorescent primers (Fig. 6D). Next, the crossover threshold (C_t_) for the miRNA and the U6 snRNA were determined from the sigmoidal amplification curves. Finally, the C_t_ values allowed us to estimate the relative-fold miRNA uptake in the mouse organs by the ΔΔC_t_ method (Fig. 6E).

*In vivo* studies with mmu-miR-298-5p were originally intended to modulate P-gp expression as recommended and suggested by our former collaborator. P-gp expression modulation by an oligonucleotide would fit well within Specific Aim #1 of the supporting NIH grant, but we eventually determined that it did not have that function [85,86,100–102]. As far as we are aware, no microRNAs are known to affect mouse P-gp expression or any related rodent transporters within the P-gp transporter superfamily. In addition, our former collaborator made miRNA sound specific, especially for P-gp transporter expression, but a single miRNA can affect many physiological functions [103]. For example, the mmu-miR-298-5p is known to target the nuclear factor (NF)-κB activator 1 (Act1) involved in cell signaling pathways and autophagy [100,104]. The miRNA is also known to regulate the expression of β-Amyloid Precursor Protein-converting Enzyme 1 [101] and regulate the expression of the insulin-like growth factor-1 β (IGF1Rβ) [102]. The miRNA also has a wide variety of theoretical targets that remain to be investigated in the TargetScanVert and miRDB databases [105,106]. Because of multiple unrelated targets for mmu-miR-298-5p, the *in vivo* mouse experiments were used to determine the mouse organ specificity by functionalized vesicle treatment and their effects on the mouse’s innate immunity.

Eventually, our laboratory wants to use functionalized vesicles to modulate *in vitro* and *in vivo* P-gp transporter expression. Luckily, siRNAs have already been shown to affect mouse P-gp expression [53,107]. Therefore, functionalized vesicles by our approach loaded with the P-gp targeting siRNA may be a practical way to investigate the P-gp transporter in the future.

### U6 snRNA normalization for miRNA uptake should be considered qualitative

MiRNA should be normalized against a stably expressing endogenous control RNA or RNAs to estimate the relative miRNA uptake within target cells. Ideal reference genes should have low expression variability and comparable expression levels since they can significantly impact miRNA quantitation [e.g., 108].

For these pilot studies, the relative miRNA uptake was normalized against U6 snRNA [109]. Normalizing against U6 snRNA demonstrated the feasibility of our approach but is not ideal for the following reasons. U6 snRNA expression can vary in mouse tissues (even within the same organ), differ significantly between genders, and be affected by a mouse’s age [108,110–113]. U6 snRNA expression also likely varies greatly between mouse strains and sub-strains [114,115]. Even the miRNA of interest can theoretically affect U6 snRNA levels. However, no significant changes in U6 snRNA expression were observed in these pilot studies by mEV and mLNP treatment (data not shown).

A table of mice used in these pilot studies is provided in Table S1 of the *Supplementary Information*. In these pilot studies, different mouse strains with different ages and genders were investigated. Because of potential differences in the U6 snRNA expression in the tissues of mice, this work’s relative miRNA uptake in the mouse organs should be considered qualitative. To roughly compare the miRNA uptake distributions between the control miRNA uptake in Fig. 7 from C57BL6/J mice and functionalized vesicles in Figs. 8 and 9 from Nu/Nu mice, the relative miRNA uptake from each organ was divided by the sum of the relative miRNA uptake in all the organs.

**Figure 7.**
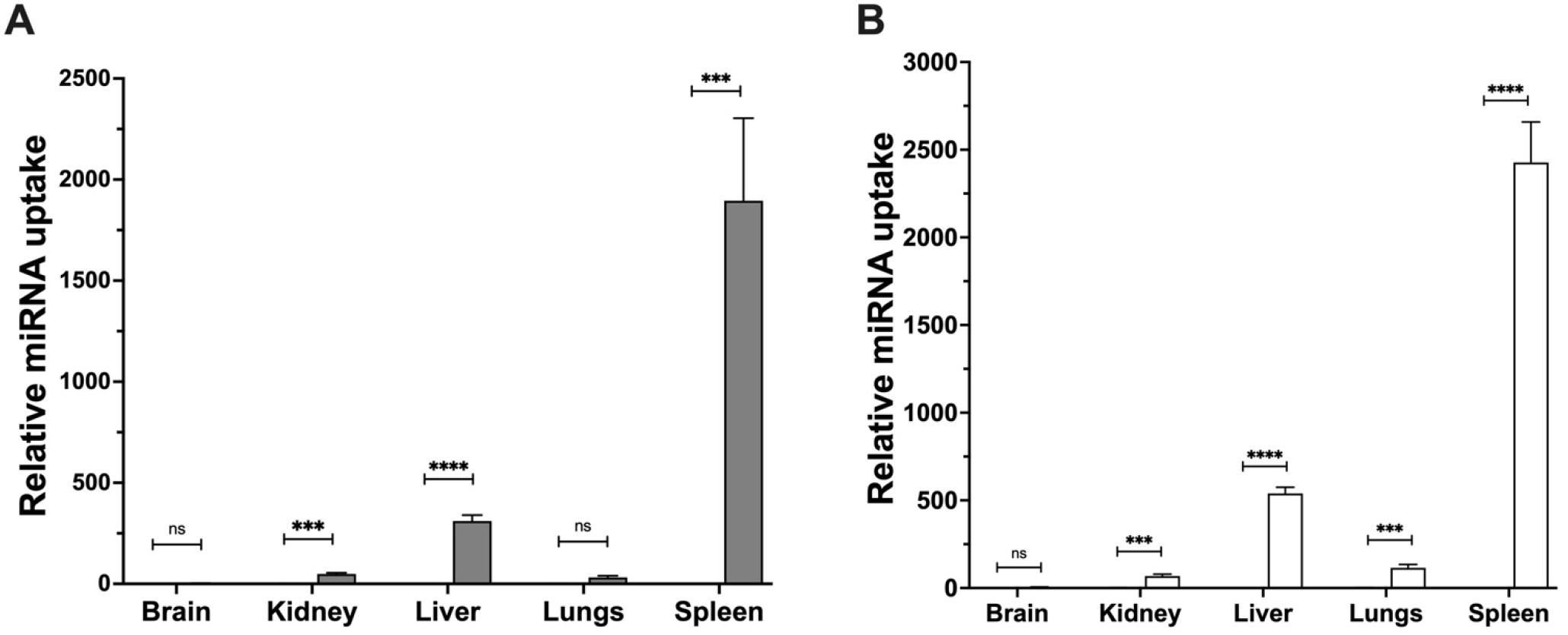
Uptake of miRNA in organs of mice treated with functionalized vesicles containing the non-targeting polyclonal green fluorescent GFP (GFP_PAB_) antibodies. The figure shows the relative-fold uptake of miRNA in the organs of young female Nu/Nu mice (n=4) treated with (A) mEVs(GFP_PAB_, mmu-miR-298) (gray) and (B) mLNPs(GFP_PAB_, mmu-miR-298) (white) versus mice that were treated with SFM only (black and barely visible in the figure). The data were normalized against the U6 snRNA and expressed as mean ± SEM. The p-values are * p ≤ 0.05, ** p ≤ 0.01, *** p ≤ 0.2 , **** p<0.0001. The dosage for each of the mice was 100 μg miRNA with an injection volume of 250-300 μl. Abbreviations: SEM, standard error of the mean.

**Figure 8.**
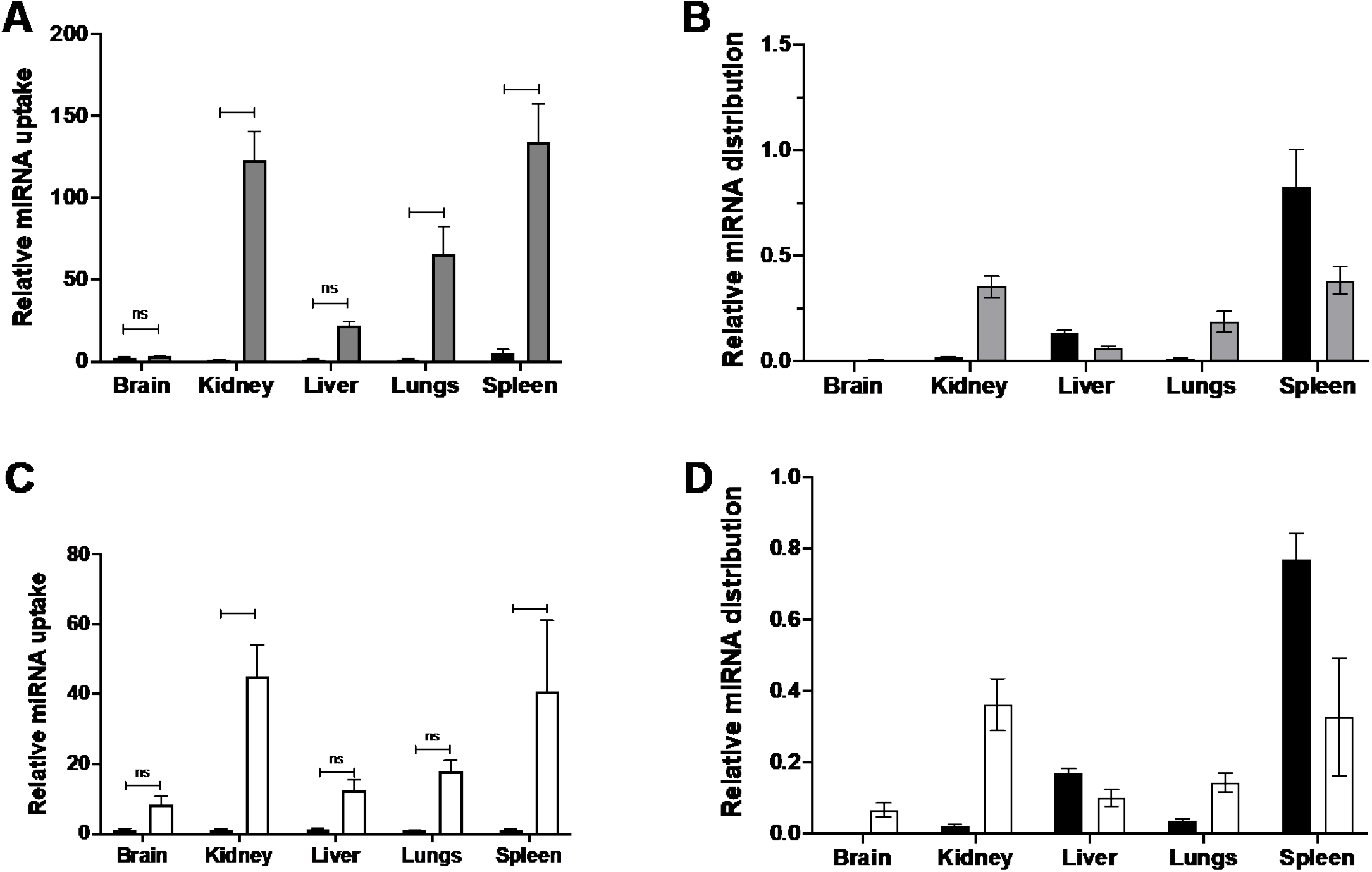
Relative mmu-miR-298 distribution by organs with the treatment of mEVs and mLNPs bioengineered with the NPHS2 antibody. The following figure shows the fold-difference of mmu-miR-298 uptake and distribution of 5-7 week old female Nu/Nu mice being treated with (A,B) mEVs when bioengineered with NPHS2 antibody (gray, n=3) (A) versus SFM only treated mice, and (B) distribution in comparison to 5-7 week old C57BL6/J female mice treated with mEV(GFP, miRNA) (black, n=4) and (C,D) mLNPs when bioengineered with NPHS2 antibody (gray)(n=6) versus (C) SFM only young female Nu/Nu mice, and (D) in comparison to when bioengineered with GFP antibody (black)(n=4) in various organs of 5-7 week old female C57BL6/J mice. All the data were normalized to the constitutive level of U6 snRNA to show uptake as well as to the total amount of miRNA delivered to show distribution and represent the mean± SEM. The p-values are * p ≤ 0.05, ** p ≤ 0.01, *** p ≤ 0.001, **** p<0.0001. The dosage for each of the mice was 100 μg with an injection volume of 250-300 μl. Abbreviations: SEM, standard error of the mean.

**Figure 9.**
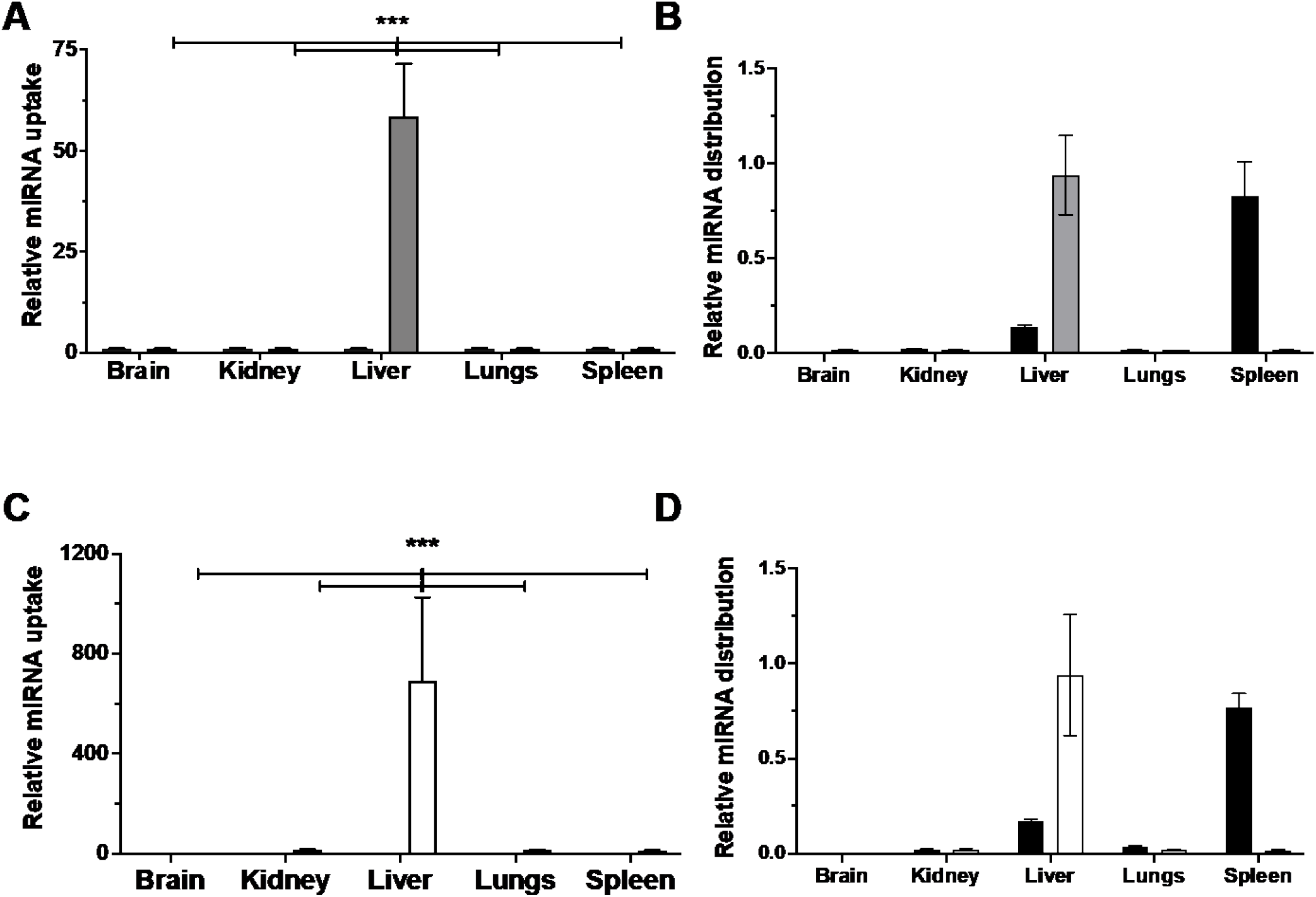
Targeting of mmu-miR-298 with the ASGR1 antibody in mice. The following figure shows the fold-difference of mmu-miR-298 uptake (A,C) and the distribution (B,D) after being treated with (A,B) mEVs when bioengineered with ASGR1 antibody (gray)(n=3) in male Nu/Nu mice aged between 15-17 weeks old versus (A) SFM only treated mice (black), and (B) versus mice treated with mEVs bioengineered with GFP antibody (black)(n=4) in female C57BL6/J mice aged 5-7 weeks old and (C,D) mLNPs when bioengineered with ASGR1 antibody (gray)(n=6) versus (C) SFM only treated female Nu/Nu mice aged 5-7 weeks old, and (D) versus mice treated with mLNPs bioengineered with GFP antibody (black) (n=4) in female C57BL6/J mice aged between 5-7 weeks old. All the data were normalized to the constitutive level of U6 snRNA to show uptake as well as to the total amount of miRNA delivered to show distribution and represent the mean ± SEM. The p-values are * p ≤ 0.05, ** p ≤ 0.01, *** p ≤ 0.001, **** p<0.0001. The dosage for each of the mice was 100 μg miRNA with an injection volume of 250-300 μl. Abbreviations: SEM, standard error of the mean.

### *In vivo* miRNA delivery by functionalized vesicles with a control antibody

As a negative control for *in-vivo* targeting ability of mEVs and mLNPs, young female C57BL6/J mice were treated with functionalized vesicles loaded with mmu-miR-298-5p and engineered with a GFP_MAB_ as a control antibody. Fig. 7 shows the uptake of mmu-miR-298 treated with modified vesicles with the GFP_MAB_ and containing mmu-miR-298 (abbreviated mEVs(GFP_MAB_, mmu-miR-298) and mLNPs(GFP_MAB_, mmu-miR-298-5p)). The highest miRNA uptake was observed in the mouse spleen after treatment with the functionalized vesicles. Uptake in the spleen is likely due to the administration route because IP administration puts the functionalized vesicle into the lymphatic system, which includes the spleen [116]. The liver had about 10-20% of the miRNA uptake than the spleen. The kidney and the lungs had only 20% of the miRNA uptake of the liver and 5% of the miRNA uptake of the spleen. Less than 1% of miRNA uptake was observed in the brain. Except for the lungs, the miRNA uptake in mice follows a very similar pattern as biotin-labeled liposomal uptake in the organs of rats [117].

### *In vivo* targeted miRNA delivery to the kidney by modified vesicles with the NPHS2 antibody

NPHS2 (a.k.a. Podocin) is a protein associated with the kidney [118]. Young female Nu/Nu mice were treated with functionalized vesicles engineered with NPHS2_PAB_ and loaded with mmu-miR-298. In Fig. 8A, the relative miRNA uptake is evenly split between the kidney and the spleen or about 125-fold higher than the U6 snRNA expression level. The mouse lung and the mouse liver had 50% and 20% of the relative miRNA uptake, respectively than the kidney and spleen. Thus, delivery of microRNA to organs outside the kidney may be due to cross-reactivity of NPHS2_PAB_ [119,120]. However, the overall miRNA uptake decreased 10-20-fold compared to Nu/Nu mice treated with mEVs(NPHS2_PAB_, mmu-miR-298) than C57BL6/J mice treated with the same dose of mEVs(GFP_MAB_, mmu-miR-298) (cf. Fig. 7A and 8A). These differences might be due to variation in U6 snRNA expression between the mouse strains, which would artifactually affect the relative miRNA uptake. Therefore, to compare the mouse strains, the relative mRNA uptake of individual organs was normalized against the total miRNA uptake of each mouse strain to produce Fig. 8B. The distribution of miRNA uptake in Fig. 8B shows that miRNA uptake by Nu/Nu mice to the spleen was 50% of C57BL6/J mice treated with mEVs but increased miRNA uptake in the kidneys. Figs. 8C shows the relative miRNA uptake of organs of mice treated with modified LNPs (abbreviated mLNPs(NPHS2_PAB_, mmu-miR-298) or mLNPs(NPHS2_PAB_, miRNA)), and it is about half the miRNA uptake as mice treated with mEVs. Still, the distribution of miRNA uptake to the organs was very similar to mEVs (Fig. 8A). The normalized distribution of mice treated with mLNPs(NPHS2_PAB_, mmu-miR-298) and mLNPs(GFP_MAB_, mmu-miR-298) in Fig. 8D was virtually identical to the distribution in Fig. 8B, implying that both types of functionalized vesicles engineered with NPHS2_PAB_ had similar targeting mechanisms in the Nu/Nu mouse strain.

### *In vivo* targeted miRNA delivery to the liver by functionalized vesicles with the ASGR1 antibody

Male and female Nu/Nu mice were treated with mEVs(ASGR1_PAB_, mmu-miR-298) and mLNPs(ASGR1_PAB_, mmu-miR-298), respectively. Figs. 9A and C show the relative miRNA uptake by various organs in adult male Nu/Nu mice treated with mEVs(ASGR1_PAB_, mmu-miR-298), and young female Nu/Nu mice were treated with mLNPs(ASGR1_PAB_, mmu-miR-298) compared to levels of mmu-miR-298 in Nu/Nu mice only treated with SFM. Almost all mmu-miR-298 (94%) was in the liver for both adult male and young female Nu/Nu mice, which shows a lot of specificity for these functionalized vesicles. In the liver lobes of adult male Nu/Nu mice, the highest concentrations of mmu-miR-298 were found in the median lobe, which is the largest, and the caudate lobe, which directly interacts with inferior vena cava in mice (Fig. S1). The relative amount of mmu-miR-298 in adult male Nu/Nu mice was about the tenth of the young female Nu/Nu mice. These differences may be an artifact of normalizing with U6 snRNA, which likely differs significantly between adult male Nu/Nu mice and young female Nu/Nu mice. Figs. 9B and 9D show the distributions of miRNA uptake between mice treated with functionalized vesicles engineered with the ASGR1_PAB_ and the GFP_MAB_. Regardless of the age or gender, the panels show that virtually all miRNA uptake by mice treated with functionalized vesicles with ASGR1_PAB_ was to the liver compared to 20% miRNA uptake by mice treated with control vesicles engineered with GFP_MAB_. The high specificity for mEVs and mLNPs that target the ASGR1 receptor of hepatocytes may be due to the endocytotic nature of the receptor [121] and access of the mesenteric circulation to the intraperitoneal cavity [122].

### Immunogenic effects of miRNA delivery by functionalized vesicles with the relatively non-specific ACE2 antibody

To explore the immunogenicity of mEVs and mLNPs *in-vivo*, a targeting vesicle was engineered with a monoclonal antibody against the membrane-associated angiotensin-converting enzyme 2 (ACE2) (ACE2_MAB_) with young female C57BL6/J mice with a functioning immune system. This enzyme is ubiquitously expressed in many organs throughout the body and plays a crucial role in controlling blood pressure [123–127]. An ACE2 mRNA and protein study in rodent organs showed the protein relatively evenly distributed between the rodent’s organs [128]. The highest ACE2 protein levels from rodents are in the small intestine, and the lowest ACE2 protein levels are in the spleen [128].

Fig. 10 shows the uptake of mmu-miR-298 and their effects on the cytokine levels after treatment of mice with miRNA-loaded mEVs and mLNPs engineered with ACE2_MAB_. Consistent with ACE2 protein levels in rodents, the miRNA uptake between the mouse organs was more evenly distributed than other targeting Abs. In Fig. 10A, from high to low miRNA uptake by mEVs(ACE2_MAB_, mmu-miR-298) treated mice versus C57BL6/J mice treated with SFM only, the miRNA uptake in mice by mEVs was ∼280 fold in the liver, ∼250 fold in the kidneys, ∼200 fold in the spleen, ∼200 fold in the lungs, ∼90 fold in the small intestine, ∼70 fold in the heart and ∼25 fold in the brain.

**Figure 10.**
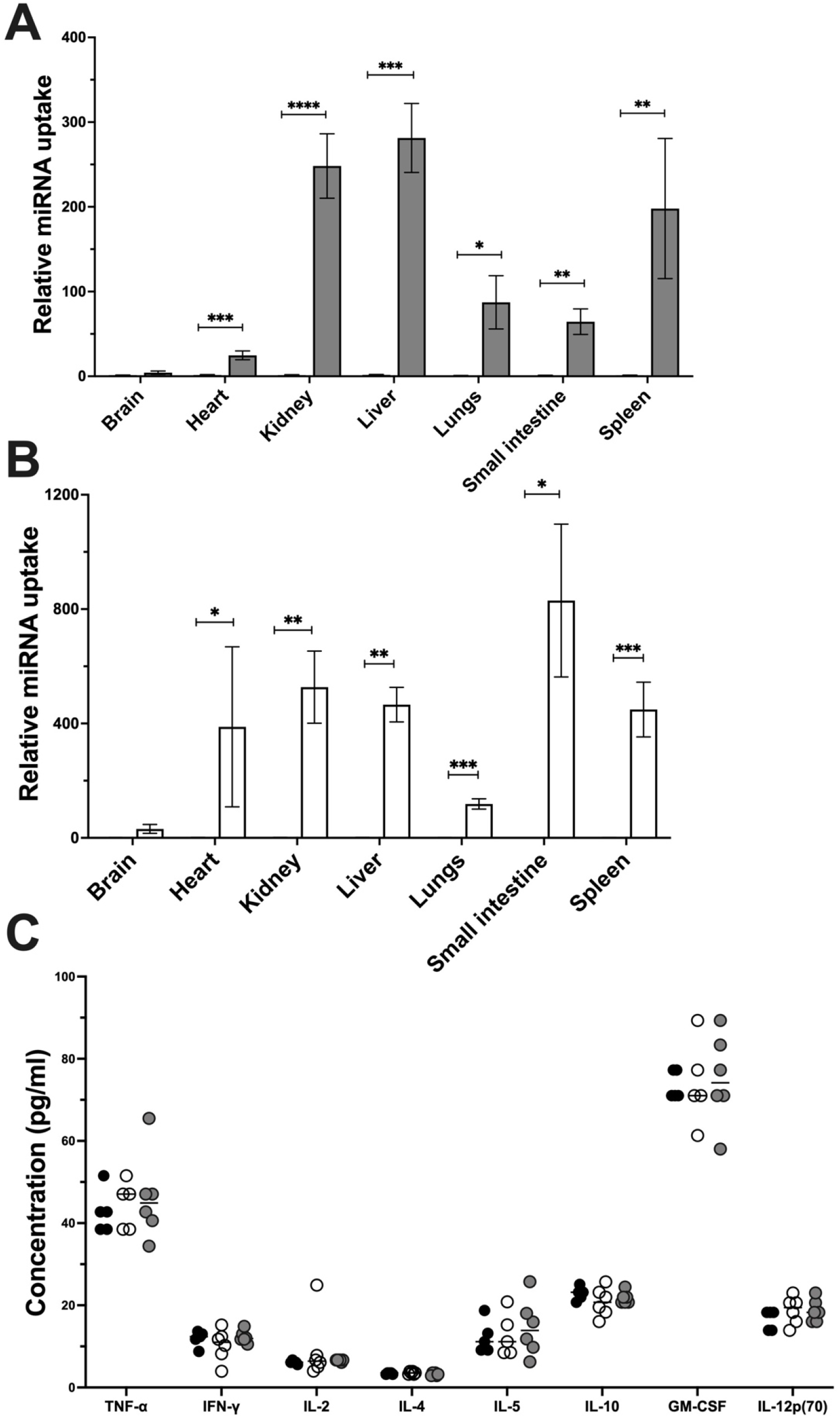
ACE2 targeting and immune reactivity of mEV(ACE2) and mLNP(ACE2) in-vivo. The relative ratio of mmu-miR-298 uptake in various organs of mice after being treated with (A) mEVs(ACE2_PAB_, miRNA) (gray) and (B) mLNP(ACE2_PAB_, miRNA) (white) versus SFM only treated female C57BL6/J mice aged 5-7 weeks (black) (n=6). The data were normalized to U6 and expressed as mean± SEM of at least three independent experiments. C) Levels of 8-major cytokine factors as obtained after conducting cytokine assay on the blood samples withdrawn from mice treated with mEVs(ACE2_PAB_, miRNA) (gray circles), mLNP(ACE2_PAB_, miRNA) (white circles), and SFM only (black circles) (n=5 or 6) 72 hours after the treatment. The line in the panel indicates a mean of 5 or 6 technical replicates from individual mice under the different treatment conditions. The p-values are * p ≤ 0.05, ** p ≤ 0.01, *** p ≤ 0.001, **** p<0.0001. The dosage for the mice was 100 μg and the injection volume was 250-300 μl. Abbreviations: SEM, standard error of the mean.

In Fig. 10B, the relative miRNA uptake was analyzed with C57BL6/J mice treated with mLNPs(ACE2_MAB_, mmu-miR-298) versus SFM only treated C57BL6/J mice. The distribution of miRNA delivery to organs by mLNPs differed from mEVs but was still well distributed between the mouse’s organs. The miRNA uptake was highest for the small intestine (∼830 fold) followed by the kidney (∼530 fold), spleen (∼400 fold), liver (∼400 fold), heart (∼388 fold), and lungs (∼100 fold). Almost no miRNA uptake was observed in the brains of these mice. Except for the spleen, these relative miRNA uptake levels are consistent with ACE2 protein levels in rodents and humans [126–128]. Since there is almost no ACE2 enzyme in the spleen [128], the relatively high miRNA uptake after treatment with functionalized vesicles is likely due to IP administration rather than specific targeting by the ACE2_MAB_.

The effects of functionalized vesicles on the mouse’s innate immunity were assessed by Bio-Plex Pro Mouse Cytokine 8-plex Assay kit. This kit tests for eight major anti- and pro-inflammatory cytokines from two types of CD4+ T helper lymphocytes [129]. Cytokine profiling has been used previously to determine the immunogenicity of EVs and LNPs [130,131]. Fig. 10C shows the cytokine levels in C57BL6/J mice treated with mEVs(ACE2_MAB_, mmu-miR-298) and mLNPs(ACE2_MAB_, mmu-miR-298) in comparison to mice treated with SFM after 72 hours. No statistically significant change in cytokine levels was observed in mice treated with either mEVs (gray circles) or mLNPs (white circles) versus the SFM only treated mice (black circles). During the 72 hour treatment period (Figs. 7-10), no physical manifestations of an innate immune response were observed, such as redness, itching, or sudden hair loss. The apparent lack of an innate immune response to the functionalized vesicles in these pilot experiments is promising, but the adaptive immunity in mice takes much longer to manifest itself [132]. Future experiments will examine the mouse’s adaptive immunity to functionalized vesicle mouse treatment over several weeks.

## Discussion

In this work, we present the “Functionalized Lipid Insertion Method” to bioengineer targeting functionalized vesicles with a surface coated with PEG-linked Abs (i.e., mEVs or mLNPs), but differs significantly from the older “Detergent-Dialysis Method” used since the early 1980s [17–24]. Both mEVs and mLNPs could efficiently deliver miRNA to the liver cancer HepG2 cell line compared to the transfection reagent. *In vivo*, the mEVs and mLNPs delivered miRNA to their targeted mouse organs. The functionalized vesicles also did not appear to significantly affect the innate immunity of mice (Fig. 10).

Even unmodified EVs can have oligonucleotide uptake efficiencies and target cells like mEVs [133,134]. Exosomes derived from the human hepatoma Huh7 cell line targeted the human embryonic kidney line over human PBMCs or a human lymphoblast cell line [133]. The miRNA uptake efficiencies for the human embryonic kidney cell line treated with exosomes were 5-200-fold higher than the other cell lines [133]. A study with exosomes derived from MDA-MB-231 cells and H-29 colon cancer cells suggested that efficient exosomal targeting relied on complementary interactions [134]. Unmodified exosomes are the simplest to produce but predicting their targeting requires analyzing the oligonucleotide uptake in various tissues and cell lines to determine specificity.

Several methods have been developed to modify the surfaces of natural vesicles to control targeting to cells and improve uptake of oligonucleotides [15,53,134]. Targeting and miRNA uptake of CD8+ T-cell-derived exosomes to effector T-cells was enhanced by having a constitutive exosomal protein bound with antibodies [15]. Unfortunately, this simple approach does not have broader applicability to exosomal targeting beyond T-cells. Another strategy to exploit constitutive exosomal proteins is to fuse them with targeting proteins by genetic engineering [135]. Exosomes were genetically engineered with a lysosomal protein (Lamp2b) and the rabies viral glycoprotein (RVG) [72]. They were able to enhance siRNA uptake efficiency in cells and overcome the blood-brain barrier (BBB) in mice [53]. Exosomes have been genetically engineered using a similar technique with fusion proteins of the lysosomal protein (Lamp2) with GFP for tracking and the HER2 affibody for targeting [136]. Exosomes have also been genetically engineered with a construct made from the vesicular stomatitis virus glycoprotein with improved loading, delivery, and tracking [137]. The development of modified exosomes by genetic engineering is limited by the complexity of the fusion protein construct and getting the construct to express in budding exosomes [135].

A highly PEGylated exosome with a relatively low surface Ab density than mEVs was developed using a two-stage process to deliver anticancer drugs to cells [16]. The modified exosomes provided only a ∼30% increase in anticancer drug uptake by a human pancreatic cancer cell line [16].

Similarly, modified and unmodified LNPs have been used to deliver miRNA previously. Cationic liposomes have been used as transfection agents to deliver various miRNAs [91,138,139]. Cationic liposomes can also form complexes with oligonucleotides such as miRNA forming lipoplexes, which have been used to deliver miRNA *in vitro* and *in vivo* [140,141]. Liposomes have been conjugated with aptamers to deliver miRNA to an ovarian cancer cell line [142]. Liposomes have been engineered with an antibody fragment to deliver siRNA and miRNA to a lung cancer cell line and tumor-bearing mice [143]. Liposomes engineered with a CD59 receptor antibody and load with an anticancer drug and miRNA increased cytotoxicity against cervical cancer cells [144]. However, liposomal engineering often relies on the older “Detergent-Dialysis Method,” which disrupts preformed vesicles and requires extensive dialysis [17–24].

Many of the pieces for industrial-scale production of the functionalized vesicles by the “Functionalized Lipid Insertion Method” are already available. Industrial methods for dialyzing and electroporation have already been around for decades [145,146]. Synthetic methods are available to produce large quantities of oligonucleotides [147]. Innovative large-scale production of monoclonal Abs is being developed from plants [148]. Currently, getting EVs is rate-limiting, but methods are on the horizon for obtaining them from abundant sources such as milk [138,139]. Liposomes have been around for decades, with different methods available for industrial production, including microfluidics [9,149,150].

Several challenges exist for implementing mEVs and mLNPs therapeutically. For long-term viability, EVs and, by extension, mEVs are typically stored at -80°C. However, month-long room temperature EV storage is possible through lyophilization with trehalose [41,151]. In contrast, lyophilized LNPs can last up to a year at 4°C or room temperature in an oxygen-free environment [152–155]. MiRNAs also have their challenges and must be stored at -80°C [156], but chemical modification can increase temperature stability considerably [157]. Another technical challenge using miRNA within functionalized vesicles is the slow leakage of miRNA from the vesicle [161].

### Future Outlook

These pilot studies demonstrated the feasibility of the “Functionalized Lipid Insertion Method” *in vitro* in HepG2 cells and *in vivo* in mice. However, additional studies will be required to unleash the full potential of the functionalized vesicles as therapeutic drug vehicles and to use them to study the biological functions of miRNA and other oligonucleotides.

Several methods are already available for LNP production, and the use of LNPs as drug delivery vehicles is well established [9,149]. However, the use of EVs as a molecular delivery vehicle is relatively new, and they have been mislabeled in the past as exosomes. EVs are actually a heterogeneous vesicle mixture [158,159]. To produce exosomes or homogenous EVs for molecular delivery, these mixtures must be appropriately processed and characterized [158,159].

To identify the best approach for *in vitro* miRNA uptake, functionalized vesicles will be engineered with different targeting moieties against a larger range of cell lines than the current pilot studies. For example, the Jurkat T-cell line could be treated with functionalized vesicles engineered with the ASGR1_PAB_ because it expresses the ASGR1 receptor [160]. An immortalized kidney cell line that expresses the NPHS2 (podocin) protein [161] could be treated with functionalized vesicles engineered with the NPHS2_PAB_. Like the in vivo experiments shown in Fig. 7, functionalized vesicles with a GFP_MAB_ can be an additional control to test the functionalized vesicle targeting ability. Investigations with different cell lines that are less characterized should include Western blots with the targeting Abs to establish appropriate targeting to the protein of interest.

Control non-coding RNA (ncRNA) used for miRNA quantitation, such as U6 snRNA, can suffer from expression instability, which can negatively impact miRNA measurements. More recent studies have emphasized the importance of carefully validating the control RNA for each cell line [108,110]. Because miRNA affects multiple mRNAs [87], a wound-healing assay of HepG2 was used to determine the functional effects of the hsa-miR-26a (Fig. 5). Investigating miRNA effects on a single protein should be monitored by a Western blot in addition to the protein’s mRNA because of potential normalization issues with control RNA.

Fig. 2B discusses the theoretical possibility of targeting secreted receptor protein ligands by our functionalized vesicles. Appropriate investigations will compare cell lines that secrete a lot of a receptor protein-ligand to ones that produce little or none of it. For example, the miRNA uptake efficiency of the MB-MDA-435 cell line that secretes a lot of the receptor protein-ligand autotaxin can be compared to the miRNA uptake efficiency of the MB-MDA-231 cell line that produces little or none of it [163–165].

The mice in a robust miRNA uptake study should be strain, gender, and age-matched. Although not shown in these pilot experiments, the degree of mEV and mLNP organ penetration should be clearly demonstrated by microscopic images of fluorescently-labeled vesicles in different parts of the mouse’s organs. Fig. 10C showed the effects of the functionalized vesicles on the innate immunity of mice and were performed as a technical replicate from mice under different treatment conditions. Future experiments should examine mice’s innate and adaptive immunity over several weeks as biological replicates where functionalized vesicle effects on mice are examined with several different mice for each treatment condition [132]. Experiments analogous to *in vitro* hsa-miR-26a delivery to HepG2 cells in Fig. 5 can be performed in mice. First, hepG2 cells can be implanted in immune system deficient nude mice (i.e., Nu/Nu) as described [166]. Then the mice can undergo treatment by functionalized vesicles loaded with hsa-miR-26 to determine their effect on HepG2 tumor size and weight.

Choosing appropriate control ncRNAs is essential for accurate miRNA quantitation. Therefore, several ncRNAs from each animal tissue or cell line should be examined for expression stability and variability. Several distinct approaches have been developed for this purpose [167–170]. Using the methods together seems the best approach to determine the optimum control ncRNA for miRNA quantitation [108]. Even after reference ncRNA genes have been identified, control RNA gene expression should be monitored in the presence of the miRNA of interest to ensure ncRNA expression is constant.

## Authors contributions

Authorship was determined using guidelines recommended by *The International Committee of Medical Journal Editors* [171–174], which requires significant and novel contributions to scholarship and significant contributions to the organization, writing, and intellectual ideas within this manuscript. The original development of the “Functionalized Lipid Insertion Method” was based on membrane protein reconstitution in liposomes previously done in Dr. Arthur G. Roberts laboratory [52,53]. Ms. Pragati Jain made significant improvements to the functionalized vesicle experimental protocol that was part of Dr. Sudeepti Kuppa’s Ph.D. thesis. Ms. Pragati Jain and Dr. Arthur G. Roberts played the largest roles in designing the experiments within the manuscript, as evidenced by their many individual meetings. They both played major roles in the writing and the organization of the manuscript. Ms. Pragati Jain selected the specific antibodies and most of the other materials that were used within the project. Dr. Arthur G. Roberts selected the fluorescent and reactive lipids used in this project and designed the original protocol for functionalized vesicles, as well as contributed to modified versions by Dr. Sudeepti Kuppa and Ms. Pragati Jain. Ms. Pragati Jain did all the data collection and almost all of the data analysis.

## Acknowledgments

We would like to thank the National Cancer Institute Grant (1R01CA204846-01A1) for the resources necessary for this research. We would like to thank the generosity of several Pharmaceutical and Biomedical Sciences (PBS) Department faculty members at the University of Georgia (UGA) for sharing their instrumentation, their cell culture hoods, their animal use protocols (AUPs), and space within the animal facilities at the Davidson Life Sciences and the Coverdell buildings at the University of Georgia, Athens, GA 30602. We greatly appreciate the help of Dr. Charnel Byrnes, who trained Ms. Pragati Jain in mammalian cell culture, PBMC vesicle isolation, and assisted her in optimizing electroporation for loading vesicles with miRNA. We want to thank Drs. Mandi M. Murph and Yao Yao for assistance with the mice experiments. We would also like to thank Dr. Balazs Rada for assisting us in getting PBMCs from human volunteers. We want to thank Dr. Sergiy Minko for providing the DLS machine for vesicle size determination from his laboratory. We would also like to thank various PBS faculty members at the UGA for giving comments and suggestions for this project. Most importantly, we would like to thank Dr. Sudeepti Kuppa, who used the original protocol that Dr. Arthur G. Roberts conceived and designed to work *in vitro* for the MDA-MB-231 cell line, which was a significant part of her Ph.D. thesis. Figs. 1, 2, 5A, and 6 were created with BioRender.com.

## Abbreviations

Ab: antibody
C_t_: cycle threshold
CMC: critical micelle concentration
EMEM: Eagle’s minimal essential medium (EMEM)
EV: extracellular vesicle
FA: fatty acid
FBS: Fetal Bovine Serum
HepG2: human liver cancer cell line
HCC: hepatocellular carcinoma
LNP: liposomal nanoparticle
miR: microRNA
miRNA: microRNA
mEV: modified extracellular vesicle
mLNP: modified liposomal nanoparticle
PBMCs: peripheral blood nuclear cells
PBS: phosphate-buffered saline.

## Supplementary Information for

**Table S1.** Mice used in these pilot studies showing the strain, gender, age and functionalized vesicle treatment given to the mice. Mice 5-7 weeks old are considered young, while mice aged 15-17 weeks are considered adults. **Abbreviations:** ASGR1_MAB_, ASGR1 receptor monoclonal antibody; GFP_MAB_, green fluorescent protein (GFP) monoclonal antibody; NPHS2_PAB_, NPHS2 or podocin polycolonal antibody; mEV, modified extracellular vesicle; mLNP, modified liposomal nanoparticle; SFM, Serum Free Media;

| Figure | Strain | Common Name | Gender | Age | # | Treatment |
| --- | --- | --- | --- | --- | --- | --- |
| 7A | C57BL6/J | C57 black 6 mouse | Female | 5-7 weeks | 4 | mEV(GFP<sub>MAB</sub>, mmu-miR-298-5p) |
| 7B | C57BL6/J | C57 black 6 mouse | Female | 5-7 weeks | 4 | mLNP(GFP<sub>MAB</sub>, mmu-miR-298-5p) |
| 7A,B | C57BL6/J | C57 black 6 mouse | Female | 5-7 weeks | 4 | SFM only |
| 8A | Nu/Nu | Immunodeficient nude mouse | Female | 5-7 weeks | 3 | mEV(NPHS2<sub>PAB</sub>, mmu-miR-298-5p) |
| 8B | Nu/Nu | Immunodeficient nude mouse | Female | 5-7 weeks | 6 | mLNP(NPHS2<sub>PAB</sub>, mmu-miR-298-5p) |
| 8A,B | Nu/Nu | Immunodeficient nude mouse | Female | 5-7 weeks | 9 | SFM only |
| 9A, S1 | Nu/Nu | Immunodeficient nude mouse | Male | 15-17 weeks | 3 | mEV(ASGR1<sub>PAB</sub>, mmu-miR-298-5p) |
| 9B | Nu/Nu | Immunodeficient nude mouse | Female | 5-7 weeks | 6 | mLNP(ASGR1<sub>PAB</sub>, mmu-miR-298-5p) |
| 9A, S1 | Nu/Nu | Immunodeficient nude mouse | Male | 15-17 weeks | 3 | SFM only |
| 9B | Nu/Nu | Immunodeficient nude mouse | Female | 5-7 weeks | 6 | SFM only |
| 10A | C57BL6/J | C57 black 6 mouse | Female | 5-7 weeks | 6 | mEV(ACE2<sub>MAB</sub>, mmu-miR-298-5p) |
| 10B | C57BL6/J | C57 black 6 mouse | Female | 5-7 weeks | 6 | mLNP(ACE2<sub>MAB</sub>, mmu-miR-298-5p) |
| 10A,B | C57BL6/J | C57 black 6 mouse | Female | 5-7 weeks | 6 | SFM only |

**Figure S1:**
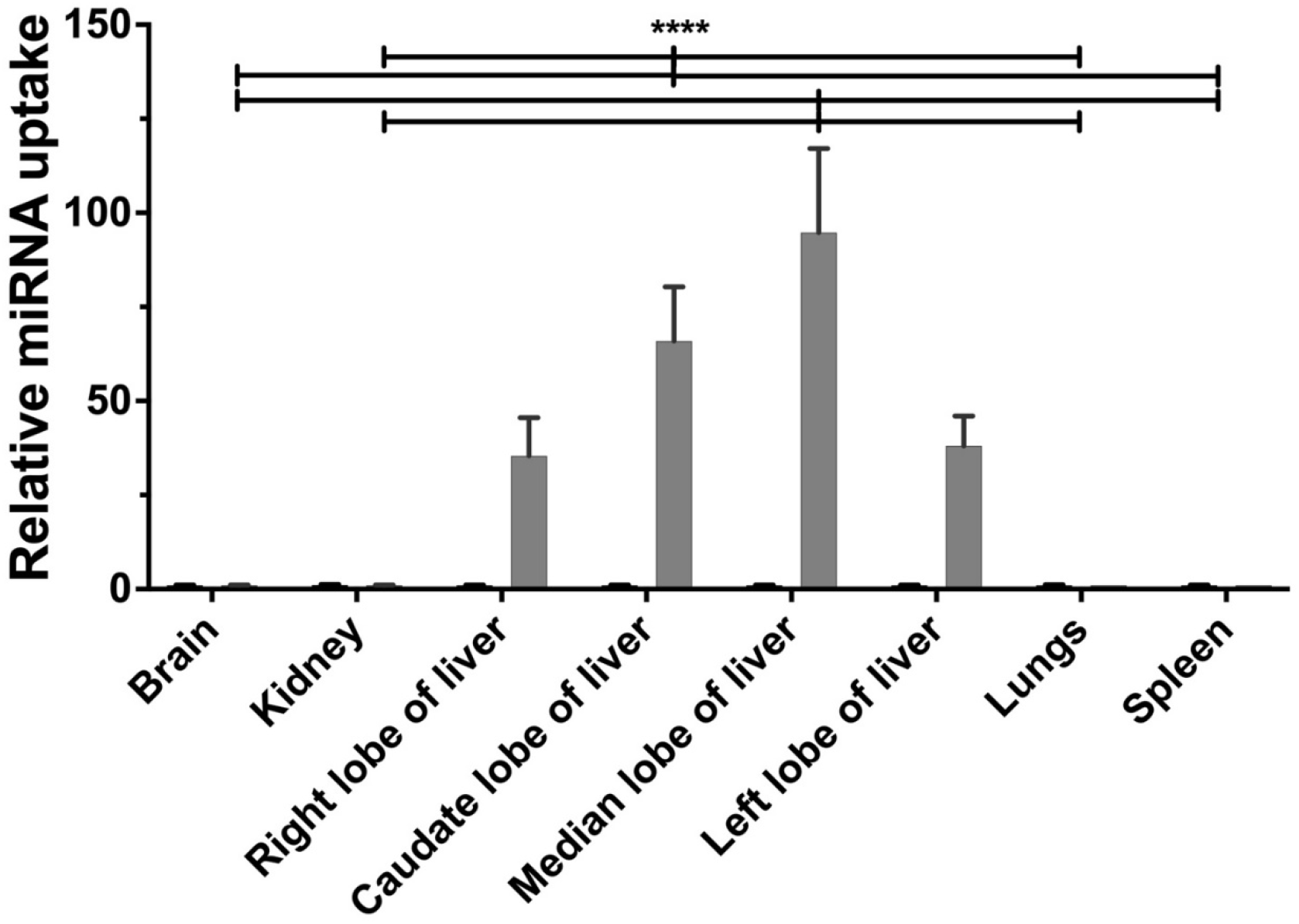
The relative miRNA uptake by different mouse liver lobes after mEV treatment of vesicles engineered with the ASGR1 receptor polyclonal antibody and loaded with mmu-miR-298. The following figure shows the relative miRNA uptake after being treated with mEVs bioengineered with the ASGR1 receptor polyclonal antibody and loaded with mmu-miR-298 (mEV(ASGR1_PAB_, mmu-miR-298)) (gray) (n=3) in adult male Nu/Nu mice versus SFM only treated adult male mice (black). All the data were normalized to the constitutive level of U6 snRNA and represented the mean ± SEM of three independent experiments. The mEV dosage given for each of the mice was 100 μg miRNA with an injection volume of 250-300 μl. All the data were normalized to the constitutive level of U6 snRNA to show uptake as well as to the total amount of miRNA delivered to show distribution and represent the mean ± SEM. The p-values are * p ≤ 0.05, ** p ≤ 0.01, *** p ≤ 0.001, **** p<0.0001. Abbreviations: SEM, standard error of the mean.

## Bibliography

[1] J.K. Patra, G. Das, L.F. Fraceto, E.V.R. Campos, M. del P. Rodriguez-Torres, L.S. Acosta-Torres, L.A. Diaz-Torres, R. Grillo, M.K. Swamy, S. Sharma, S. Habtemariam, H.-S. Shin, Nano based drug delivery systems: recent developments and future prospects, Journal of Nanobiotechnology. 16 (2018) 71. https://doi.org/10.1186/s12951-018-0392-8.

[2] D. Das, N. Maity, A.H. V, Nanotechnology: a revolution in targeted drug delivery, International Journal of Basic & Clinical Pharmacology. 6 (2017) 2766–2773. https://doi.org/10.18203/2319-2003.ijbcp20175200.

[3] B.S. Zolnik, Á. González-Fernández, N. Sadrieh, M.A. Dobrovolskaia, Nanoparticles and the Immune System, Endocrinology. 151 (2010) 458–465. https://doi.org/10.1210/en.2009-1082.

[4] S. Su, P.M. Kang, Systemic Review of Biodegradable Nanomaterials in Nanomedicine, Nanomaterials (Basel). 10 (2020). https://doi.org/10.3390/nano10040656.

[5] X. Li, L. Wang, Y. Fan, Q. Feng, F. Cui, Biocompatibility and Toxicity of Nanoparticles and Nanotubes, Journal of Nanomaterials. 2012 (2012) e548389. https://doi.org/10.1155/2012/548389.

[6] F. Alexis, E. Pridgen, L.K. Molnar, O.C. Farokhzad, Factors Affecting the Clearance and Biodistribution of Polymeric Nanoparticles, Mol Pharm. 5 (2008) 505–515. https://doi.org/10.1021/mp800051m.

[7] S.T. Jahan, S.M.A. Sadat, M. Walliser, A. Haddadi, Targeted Therapeutic Nanoparticles: An Immense Promise to Fight against Cancer, Journal of Drug Delivery. 2017 (2017) e9090325. https://doi.org/10.1155/2017/9090325.

[8] X. Luan, K. Sansanaphongpricha, I. Myers, H. Chen, H. Yuan, D. Sun, Engineering exosomes as refined biological nanoplatforms for drug delivery, Acta Pharmacologica Sinica. 38 (2017) 754–763. https://doi.org/10.1038/aps.2017.12.

[9] A. Akbarzadeh, R. Rezaei-Sadabady, S. Davaran, S.W. Joo, N. Zarghami, Y. Hanifehpour, M. Samiei, M. Kouhi, K. Nejati-Koshki, Liposome: classification, preparation, and applications, Nanoscale Res Lett. 8 (2013) 102. https://doi.org/10.1186/1556-276X-8-102.

[10] E.M. Veziroglu, G.I. Mias, Characterizing Extracellular Vesicles and Their Diverse RNA Contents, Front Genet. 11 (2020) 700. https://doi.org/10.3389/fgene.2020.00700.

[11] R. Kalluri, V.S. LeBleu, The biology, function, and biomedical applications of exosomes, Science. 367 (2020). https://doi.org/10.1126/science.aau6977.

[12] J. Qin, Q. Xu, Functions and application of exosomes, Acta Pol Pharm. 71 (2014) 537–43.

[13] R.R. Sawant, V.P. Torchilin, Challenges in Development of Targeted Liposomal Therapeutics, AAPS J. 14 (2012) 303–315. https://doi.org/10.1208/s12248-012-9330-0.

[14] A. Samad, Y. Sultana, M. Aqil, Liposomal drug delivery systems: an update review, Curr Drug Deliv. 4 (2007) 297–305. https://doi.org/10.2174/156720107782151269.

[15] K. Bryniarski, W. Ptak, A. Jayakumar, K. Püllmann, M.J. Caplan, A. Chairoungdua, J. Lu, B.D. Adams, E. Sikora, K. Nazimek, S. Marquez, S.H. Kleinstein, P. Sangwung, Y. Iwakiri, E. Delgato, F. Redegeld, B.R. Blokhuis, J. Wojcikowski, A.W. Daniel, T.G. Kormelink, P.W. Askenase, Antigen-specific, antibody-coated, exosome-like nanovesicles deliver suppressor T-cell microRNA-150 to effector T cells to inhibit contact sensitivity, Journal of Allergy and Clinical Immunology. 132 (2013) 170–181.e9. https://doi.org/10.1016/j.jaci.2013.04.048.

[16] Y. Si, S. Kim, E. Zhang, Y. Tang, R. Jaskula-Sztul, J.M. Markert, H. Chen, L. Zhou, X. (Margaret) Liu, Targeted Exosomes for Drug Delivery: Biomanufacturing, Surface Tagging, and Validation, Biotechnology Journal. 15 (2020) 1900163. https://doi.org/10.1002/biot.201900163.

[17] A. Huang, L. Huang, S.J. Kennel, Monoclonal antibody covalently coupled with fatty acid. A reagent for in vitro liposome targeting., Journal of Biological Chemistry. 255 (1980) 8015–8018. https://doi.org/10.1016/S0021-9258(19)70595-X.

[18] J.L. Rigaud, D. Levy, Reconstitution of membrane proteins into liposomes, Methods in Enzymology. 372 (2003) 65–86. https://doi.org/10.1016/S0076-6879(03)72004-7.

[19] L.E. Westerman, P.E. Jensen, Liposomes Composed of Reconstituted Membranes for Induction of Tumor-Specific Immunity, in: Methods in Enzymology, Academic Press, 2003: pp. 118–127. https://doi.org/10.1016/S0076-6879(03)73008-0.

[20] A. Huang, S.J. Kennel, L. Huang, Immunoliposome labeling: A sensitive and specific method for cell surface labeling, Journal of Immunological Methods. 46 (1981) 141–151. https://doi.org/10.1016/0022-1759(81)90131-9.

[21] H. Alpes, K. Allmann, H. Plattner, J. Reichert, R. Rick, S. Schulz, Formation of large unilamellar vesicles using alkyl maltoside detergents, Biochimica et Biophysica Acta (BBA) - Biomembranes. 862 (1986) 294–302. https://doi.org/10.1016/0005-2736(86)90231-2.

[22] L. Huang, United States Patent: 4957735 - Target-sensitive immunoliposomes-preparation and characterization, 4957735, 1990. https://patft.uspto.gov/netacgi/nph-Parser?Sect1=PTO1&Sect2=HITOFF&d=PALL&p=1&u=%2Fnetahtml%2FPTO%2Fsrchnum.htm&r=1&f=G&l=50&s1=4,957,735.PN.&OS=PN/4,957,735&RS=PN/4,957,735 (accessed August 31, 2021).

[23] R.J.Y. Ho, B.T. Rouse, L. Huang, Target-sensitive immunoliposomes: preparation and characterization, Biochemistry. 25 (1986) 5500–5506. https://doi.org/10.1021/bi00367a023.

[24] M. Micklus, N. Greig, S. Rapoport, Targeting of liposomes to the blood-brain barrier, US20020025313A1, 2002. https://patents.google.com/patent/US20020025313A1/en (accessed August 21, 2021).

[25] Y. Cheng, Q. Zeng, Q. Han, W. Xia, Effect of pH, temperature and freezing-thawing on quantity changes and cellular uptake of exosomes, Protein Cell. 10 (2019) 295–299. https://doi.org/10.1007/s13238-018-0529-4.

[26] H. Ma, C. Ó’Fágáin, R. O’Kennedy, Antibody stability: A key to performance - Analysis, influences and improvement, Biochimie. 177 (2020) 213–225. https://doi.org/10.1016/j.biochi.2020.08.019.

[27] C.I.E. Smith, R. Zain, Therapeutic Oligonucleotides: State of the Art, Annual Review of Pharmacology and Toxicology. 59 (2019) 605–630. https://doi.org/10.1146/annurev-pharmtox-010818-021050.

[28] T.C. Roberts, R. Langer, M.J.A. Wood, Advances in oligonucleotide drug delivery, Nat Rev Drug Discov. 19 (2020) 673–694. https://doi.org/10.1038/s41573-020-0075-7.

[29] K.E. Lundin, O. Gissberg, C.I.E. Smith, R. Zain, Chemical Development of Therapeutic Oligonucleotides, in: O. Gissberg, R. Zain, K.E. Lundin (Eds.), Oligonucleotide-Based Therapies: Methods and Protocols, Springer, New York, NY, 2019: pp. 3–16. https://doi.org/10.1007/978-1-4939-9670-4_1.

[30] K. Tatiparti, S. Sau, S.K. Kashaw, A.K. Iyer, siRNA Delivery Strategies: A Comprehensive Review of Recent Developments, Nanomaterials. 7 (2017) 77. https://doi.org/10.3390/nano7040077.

[31] K. Gavrilov, W.M. Saltzman, Therapeutic siRNA: Principles, Challenges, and Strategies, Yale J Biol Med. 85 (2012) 187–200.

[32] K. Dhuri, C. Bechtold, E. Quijano, H. Pham, A. Gupta, A. Vikram, R. Bahal, Antisense Oligonucleotides: An Emerging Area in Drug Discovery and Development, Journal of Clinical Medicine. 9 (2020) 2004. https://doi.org/10.3390/jcm9062004.

[33] M. Gagliardi, A.T. Ashizawa, The Challenges and Strategies of Antisense Oligonucleotide Drug Delivery, Biomedicines. 9 (2021) 433. https://doi.org/10.3390/biomedicines9040433.

[34] R. Rupaimoole, F.J. Slack, MicroRNA therapeutics: towards a new era for the management of cancer and other diseases, Nat Rev Drug Discov. 16 (2017) 203–222. https://doi.org/10.1038/nrd.2016.246.

[35] R.C. Wilson, J.A. Doudna, Molecular Mechanisms of RNA Interference, Annual Review of Biophysics. 42 (2013) 217–239. https://doi.org/10.1146/annurev-biophys-083012-130404.

[36] M.M. Zhang, R. Bahal, T.P. Rasmussen, J.E. Manautou, X. Zhong, The growth of siRNA-based therapeutics: Updated clinical studies, Biochemical Pharmacology. 189 (2021) 114432. https://doi.org/10.1016/j.bcp.2021.114432.

[37] S. Akhtar, M.D. Hughes, A. Khan, M. Bibby, M. Hussain, Q. Nawaz, J. Double, P. Sayyed, The delivery of antisense therapeutics, Advanced Drug Delivery Reviews. 44 (2000) 3–21. https://doi.org/10.1016/S0169-409X(00)00080-6.

[38] J. O’Brien, H. Hayder, Y. Zayed, C. Peng, Overview of MicroRNA Biogenesis, Mechanisms of Actions, and Circulation, Front. Endocrinol. 9 (2018). https://doi.org/10.3389/fendo.2018.00402.

[39] C.E. Condrat, D.C. Thompson, M.G. Barbu, O.L. Bugnar, A. Boboc, D. Cretoiu, N. Suciu, S.M. Cretoiu, S.C. Voinea, miRNAs as Biomarkers in Disease: Latest Findings Regarding Their Role in Diagnosis and Prognosis, Cells. 9 (2020). https://doi.org/10.3390/cells9020276.

[40] W. Si, J. Shen, H. Zheng, W. Fan, The role and mechanisms of action of microRNAs in cancer drug resistance, Clinical Epigenetics. 11 (2019) 25. https://doi.org/10.1186/s13148-018-0587-8.

[41] A. Jeyaram, S.M. Jay, Preservation and Storage Stability of Extracellular Vesicles for Therapeutic Applications, AAPS J. 20 (2017) 1. https://doi.org/10.1208/s12248-017-0160-y.

[42] S.K. Panda, B. Ravindran, F. Histopaque, Isolation of human PBMCs, Bio-Protocol. 3 (2013) e323.

[43] G.E. Moore, R.E. Gerner, H.A. Franklin, Culture of Normal Human Leukocytes, JAMA. 199 (1967) 519–524. https://doi.org/10.1001/jama.1967.03120080053007.

[44] B. Mui, L. Chow, M.J. Hope, Extrusion technique to generate liposomes of defined size, Methods in Enzymology. 367 (2003) 3–14. https://doi.org/10.1016/S0076-6879(03)67001-1.

[45] K.V. Ledwitch, R.W. Barnes, A.G. Roberts, Unravelling the complex drug-drug interactions of the cardiovascular drugs, verapamil and digoxin, with P-glycoprotein, Biosci. Rep. 36 (2016) 1–14. https://doi.org/10.1042/BSR20150317.

[46] J.P. Kampf, D. Cupp, A.M. Kleinfeld, Different Mechanisms of Free Fatty Acid Flip-Flop and Dissociation Revealed by Temperature and Molecular Species Dependence of Transport across Lipid Vesicles*, Journal of Biological Chemistry. 281 (2006) 21566–21574. https://doi.org/10.1074/jbc.M602067200.

[47] S. Hupfeld, A.M. Holsæter, M. Skar, C.B. Frantzen, M. Brandl, Liposome Size Analysis by Dynamic/Static Light Scattering upon Size Exclusion-/Field Flow-Fractionation, Journal of Nanoscience and Nanotechnology. 6 (2006) 3025–3031. https://doi.org/10.1166/jnn.2006.454.

[48] G. Liang, S. Kan, Y. Zhu, S. Feng, W. Feng, S. Gao, Engineered exosome-mediated delivery of functionally active miR-26a and its enhanced suppression effect in HepG2 cells, Int J Nanomedicine. 13 (2018) 585–599. https://doi.org/10.2147/IJN.S154458.

[49] O. López, A. de la Maza, L. Coderch, C. López-Iglesias, E. Wehrli, J.L. Parra, Direct formation of mixed micelles in the solubilization of phospholipid liposomes by Triton X-100, FEBS Letters. 426 (1998) 314–318. https://doi.org/10.1016/S0014-5793(98)00363-9.

[50] V. Palmieri, D. Lucchetti, I. Gatto, A. Maiorana, M. Marcantoni, G. Maulucci, M. Papi, R. Pola, M. De Spirito, A. Sgambato, Dynamic light scattering for the characterization and counting of extracellular vesicles: a powerful noninvasive tool, J Nanopart Res. 16 (2014) 2583. https://doi.org/10.1007/s11051-014-2583-z.

[51] R. Szatanek, M. Baj-Krzyworzeka, J. Zimoch, M. Lekka, M. Siedlar, J. Baran, The Methods of Choice for Extracellular Vesicles (EVs) Characterization, Int J Mol Sci. 18 (2017). https://doi.org/10.3390/ijms18061153.

[52] I. Makra, P. Terejánszky, R.E. Gyurcsányi, A method based on light scattering to estimate the concentration of virus particles without the need for virus particle standards, MethodsX. 2 (2015) 91–99. https://doi.org/10.1016/j.mex.2015.02.003.

[53] L. Alvarez-Erviti, Y. Seow, H. Yin, C. Betts, S. Lakhal, M.J.A. Wood, Delivery of siRNA to the mouse brain by systemic injection of targeted exosomes, Nature Biotechnology. 29 (2011) 341–345. https://doi.org/10.1038/nbt.1807.

[54] S. Pirkmajer, A.V. Chibalin, Serum starvation: caveat emptor, American Journal of Physiology-Cell Physiology. 301 (2011) C272–C279. https://doi.org/10.1152/ajpcell.00091.2011.

[55] M.W. Pfaffl, Relative quantitation, in: M.T. Dorak (Ed.), Real-Time PCR, 1st edition, Taylor & Francis, 2007.

[56] A.L. Didychuk, S.E. Butcher, D.A. Brow, The life of U6 small nuclear RNA, from cradle to grave, RNA. 24 (2018) 437–460. https://doi.org/10.1261/rna.065136.117.

[57] R.L. Causin, D. Pessôa-Pereira, K.C.B. Souza, A.F. Evangelista, R.M.V. Reis, J.H.T.G. Fregnani, M.M.C. Marques, Identification and performance evaluation of housekeeping genes for microRNA expression normalization by reverse transcription-quantitative PCR using liquid-based cervical cytology samples, Oncol Lett. 18 (2019) 4753–4761. https://doi.org/10.3892/ol.2019.10824.

[58] T.D. Schmittgen, K.J. Livak, Analyzing real-time PCR data by the comparative C T method, Nature Protocols. 3 (2008) 1101–1108. https://doi.org/10.1038/nprot.2008.73.

[59] C. Km, S. Ml, R. An, K. La, Carbon Dioxide for Euthanasia: Concerns Regarding Pain and Distress, With Special Reference to Mice and Rats, Laboratory Animals. 39 (2005). https://doi.org/10.1258/0023677053739747.

[60] TRIzol Products - US, (n.d.). https://www.thermofisher.com/us/en/home/brands/product-brand/trizol.html (accessed May 16, 2020).

[61] Bio-Plex Pro^TM^ Mouse Cytokine, Chemokine, and Growth Factor Assays | Life Science Research | Bio-Rad, (n.d.). https://www.bio-rad.com/en-us/category/bio-plex-pro-mouse-cytokine-chemokine-growth-factor-assays?ID=37dd4c58-2aa7-4991-a7cb-5c51024959c6 (accessed December 11, 2020).

[62] L.A. Mulcahy, R.C. Pink, D.R.F. Carter, Routes and mechanisms of extracellular vesicle uptake, J Extracell Vesicles. 3 (2014). https://doi.org/10.3402/jev.v3.24641.

[63] G. Costaguta, G.S. Payne, Overview of Protein Trafficking Mechanisms, in: N. Segev (Ed.), Trafficking Inside Cells: Pathways, Mechanisms and Regulation, Springer, New York, NY, 2009: pp. 105–118. https://doi.org/10.1007/978-0-387-93877-6_6.

[64] A. Perrakis, W.H. Moolenaar, Autotaxin: structure-function and signaling, J. Lipid Res. 55 (2014) 1010–1018. https://doi.org/10.1194/jlr.R046391.

[65] J. Stetefeld, S.A. McKenna, T.R. Patel, Dynamic light scattering: a practical guide and applications in biomedical sciences, Biophys Rev. 8 (2016) 409–427. https://doi.org/10.1007/s12551-016-0218-6.

[66] P.M. Carvalho, M.R. Felício, N.C. Santos, S. Gonçalves, M.M. Domingues, Application of Light Scattering Techniques to Nanoparticle Characterization and Development, Frontiers in Chemistry. 6 (2018) 237. https://doi.org/10.3389/fchem.2018.00237.

[67] K.L. Linegar, A.E. Adeniran, A.F. Kostko, M.A. Anisimov, Hydrodynamic radius of polyethylene glycol in solution obtained by dynamic light scattering, Colloid J. 72 (2010) 279–281. https://doi.org/10.1134/S1061933X10020195.

[68] S. Niimi, [Determination of the particle size and relative light scattering intensity of aggregates of human IgG and humanized monoclonal antibody product induced by various stress using dynamic light scattering], Kokuritsu Iyakuhin Shokuhin Eisei Kenkyusho Hokoku. (2011) 55–60.

[69] S. Hu, L. Musante, D. Tataruch, X. Xu, O. Kretz, M. Henry, P. Meleady, H. Luo, H. Zou, Y. Jiang, H. Holthofer, Purification and Identification of Membrane Proteins from Urinary Extracellular Vesicles using Triton X-114 Phase Partitioning, J. Proteome Res. 17 (2018) 86–96. https://doi.org/10.1021/acs.jproteome.7b00386.

[70] M.T. Donato, L. Tolosa, M.J. Gómez-Lechón, Culture and Functional Characterization of Human Hepatoma HepG2 Cells, Methods Mol. Biol. 1250 (2015) 77–93. https://doi.org/10.1007/978-1-4939-2074-7_5.

[71] S. Wilkening, F. Stahl, A. Bader, Comparison of Primary Human Hepatocytes and Hepatoma Cell Line Hepg2 with Regard to Their Biotransformation Properties, Drug Metab Dispos. 31 (2003) 1035–1042. https://doi.org/10.1124/dmd.31.8.1035.

[72] G.-H. Qiu, X. Xie, F. Xu, X. Shi, Y. Wang, L. Deng, Distinctive pharmacological differences between liver cancer cell lines HepG2 and Hep3B, Cytotechnology. 67 (2015) 1–12. https://doi.org/10.1007/s10616-014-9761-9.

[73] D. Witzigmann, L. Quagliata, S.H. Schenk, C. Quintavalle, L.M. Terracciano, J. Huwyler, Variable asialoglycoprotein receptor 1 expression in liver disease: Implications for therapeutic intervention, Hepatology Research. 46 (2016) 686–696. https://doi.org/10.1111/hepr.12599.

[74] S. Das, P. Kudale, P. Dandekar, P.V. Devarajan, Asialoglycoprotein Receptor and Targeting Strategies, in: P.V. Devarajan, P. Dandekar, A.A. D’Souza (Eds.), Targeted Intracellular Drug Delivery by Receptor Mediated Endocytosis, Springer International Publishing, Cham, 2019: pp. 353–381. https://doi.org/10.1007/978-3-030-29168-6_12.

[75] Z. Shen, W. Wei, H. Tanaka, K. Kohama, G. Ma, T. Dobashi, Y. Maki, H. Wang, J. Bi, S. Dai, A galactosamine-mediated drug delivery carrier for targeted liver cancer therapy, Pharmacological Research. 64 (2011) 410–419. https://doi.org/10.1016/j.phrs.2011.06.015.

[76] S. Pranatharthiharan, M.D. Patel, V.C. Malshe, V. Pujari, A. Gorakshakar, M. Madkaikar, K. Ghosh, P.V. Devarajan, Asialoglycoprotein receptor targeted delivery of doxorubicin nanoparticles for hepatocellular carcinoma, Drug Delivery. 24 (2017) 20–29. https://doi.org/10.1080/10717544.2016.1225856.

[77] M. Wei, Y. Xu, Q. Zou, L. Tu, C. Tang, T. Xu, L. Deng, C. Wu, Hepatocellular carcinoma targeting effect of PEGylated liposomes modified with lactoferrin, European Journal of Pharmaceutical Sciences. 46 (2012) 131–141. https://doi.org/10.1016/j.ejps.2012.02.007.

[78] A.A. D’Souza, P.V. Devarajan, Asialoglycoprotein receptor mediated hepatocyte targeting — Strategies and applications, Journal of Controlled Release. 203 (2015) 126–139. https://doi.org/10.1016/j.jconrel.2015.02.022.

[79] A.G. Roberts, The Structure and Mechanism of Drug Transporters, in: S. Nagar, U.A. Argikar, D. Tweedie (Eds.), Enzyme Kinetics in Drug Metabolism: Fundamentals and Applications, Springer US, New York, NY, 2021: pp. 193–234. https://doi.org/10.1007/978-1-0716-1554-6_8.

[80] J. Bentz, M.P. O’Connor, D. Bednarczyk, J. Coleman, C. Lee, J. Palm, Y.A. Pak, E.S. Perloff, E. Reyner, P. Balimane, M. Brännström, X. Chu, C. Funk, A. Guo, I. Hanna, K. Herédi-Szabó, K. Hillgren, L. Li, E. Hollnack-Pusch, M. Jamei, X. Lin, A.K. Mason, S. Neuhoff, A. Patel, L. Podila, E. Plise, G. Rajaraman, L. Salphati, E. Sands, M.E. Taub, J.-S. Taur, D. Weitz, H.M. Wortelboer, C.Q. Xia, G. Xiao, J. Yabut, T. Yamagata, L. Zhang, H. Ellens, Variability in P-Glycoprotein Inhibitory Potency (IC50) Using Various in Vitro Experimental Systems: Implications for Universal Digoxin Drug-Drug Interaction Risk Assessment Decision Criteria, Drug Metab Dispos. 41 (2013) 1347–1366. https://doi.org/10.1124/dmd.112.050500.

[81] L.-B. Goh, K.J. Spears, D. Yao, A. Ayrton, P. Morgan, C. Roland Wolf, T. Friedberg, Endogenous drug transporters in in vitro and in vivo models for the prediction of drug disposition in man, Biochemical Pharmacology. 64 (2002) 1569–1578. https://doi.org/10.1016/S0006-2952(02)01355-2.

[82] J.D. Dukes, P. Whitley, A.D. Chalmers, The MDCK variety pack: choosing the right strain, BMC Cell Biol. 12 (2011) 43. https://doi.org/10.1186/1471-2121-12-43.

[83] J. Li, Y. Wang, I.J. Hidalgo, Kinetic analysis of human and canine P-glycoprotein-mediated drug transport in MDR1–MDCK cell model: Approaches to reduce false-negative substrate classification, Journal of Pharmaceutical Sciences. 102 (2013) 3436–3446. https://doi.org/10.1002/jps.23523.

[84] P. Anderle, E. Niederer, W. Rubas, C. Hilgendorf, H. Spahn-Langguth, H. Wunderli-Allenspach, H.P. Merkle, P. Langguth, P-Glycoprotein (P-gp) mediated efflux in Caco-2 cell monolayers: the influence of culturing conditions and drug exposure on P-gp expression levels, J Pharm Sci. 87 (1998) 757–762. https://doi.org/10.1021/js970372e.

[85] L. Bao, S. Hazari, S. Mehra, D. Kaushal, K. Moroz, S. Dash, Increased expression of P-glycoprotein and doxorubicin chemoresistance of metastatic breast cancer is regulated by miR-298, Am. J. Pathol. 180 (2012) 2490–2503. https://doi.org/10.1016/j.ajpath.2012.02.024.

[86] Y. Xie, Y. Shao, X. Deng, M. Wang, Y. Chen, MicroRNA-298 Reverses Multidrug Resistance to Antiepileptic Drugs by Suppressing MDR1/P-gp Expression in vitro, Front Neurosci. 12 (2018). https://doi.org/10.3389/fnins.2018.00602.

[87] A. Kozomara, M. Birgaoanu, S. Griffiths-Jones, miRBase: from microRNA sequences to function, Nucleic Acids Res. 47 (2019) D155–D162. https://doi.org/10.1093/nar/gky1141.

[88] A. Marco, J.I. MacPherson, M. Ronshaugen, S. Griffiths-Jones, MicroRNAs from the same precursor have different targeting properties, Silence. 3 (2012) 8. https://doi.org/10.1186/1758-907X-3-8.

[89] P.E. Marques, S. Nyegaard, R.F. Collins, F. Troise, S.A. Freeman, W.S. Trimble, S. Grinstein, Multimerization and Retention of the Scavenger Receptor SR-B1 in the Plasma Membrane, Developmental Cell. 50 (2019) 283–295.e5. https://doi.org/10.1016/j.devcel.2019.05.026.

[90] D. Sahoo, Y.F. Darlington, D. Pop, D.L. Williams, M.A. Connelly, Scavenger receptor class B Type I (SR-BI) assembles into detergent-sensitive dimers and tetramers, Biochimica et Biophysica Acta (BBA) - Molecular and Cell Biology of Lipids. 1771 (2007) 807–817. https://doi.org/10.1016/j.bbalip.2006.03.003.

[91] X. Cui, K. Song, X. Lu, W. Feng, W. Di, Liposomal Delivery of MicroRNA-7 Targeting EGFR to Inhibit the Growth, Invasion, and Migration of Ovarian Cancer, ACS Omega. (2021). https://doi.org/10.1021/acsomega.1c00992.

[92] M.T.D. Martino, V. Campani, G. Misso, M.E.G. Cantafio, A. Gullà, U. Foresta, P.H. Guzzi, M. Castellano, A. Grimaldi, V. Gigantino, R. Franco, S. Lusa, M. Cannataro, P. Tagliaferri, G.D. Rosa, P. Tassone, M. Caraglia, In Vivo Activity of MiR-34a Mimics Delivered by Stable Nucleic Acid Lipid Particles (SNALPs) against Multiple Myeloma, PLOS ONE. 9 (2014) e90005. https://doi.org/10.1371/journal.pone.0090005.

[93] A. Stamm, K. Reimers, S. Strauß, P. Vogt, T. Scheper, I. Pepelanova, In vitro wound healing assays – state of the art, BioNanoMaterials. 17 (2016) 79–87. https://doi.org/10.1515/bnm-2016-0002.

[94] J.E.N. Jonkman, J.A. Cathcart, F. Xu, M.E. Bartolini, J.E. Amon, K.M. Stevens, P. Colarusso, An introduction to the wound healing assay using live-cell microscopy, Cell Adhesion & Migration. 8 (2014) 440–451. https://doi.org/10.4161/cam.36224.

[95] M.J. Haney, Y. Zhao, J. Fay, H. Duhyeong, M. Wang, H. Wang, Z. Li, Y.Z. Lee, M.K. Karuppan, N. El-Hage, A.V. Kabanov, E.V. Batrakova, Genetically modified macrophages accomplish targeted gene delivery to the inflamed brain in transgenic Parkin Q311X(A) mice: importance of administration routes, Scientific Reports. 10 (2020) 11818. https://doi.org/10.1038/s41598-020-68874-7.

[96] O.P.B. Wiklander, J.Z. Nordin, A. O’Loughlin, Y. Gustafsson, G. Corso, I. Mäger, P. Vader, Y. Lee, H. Sork, Y. Seow, N. Heldring, L. Alvarez-Erviti, C.E. Smith, K. Le Blanc, P. Macchiarini, P. Jungebluth, M.J.A. Wood, S.E. Andaloussi, Extracellular vesicle in vivo biodistribution is determined by cell source, route of administration and targeting, J Extracell Vesicles. 4 (2015) 10.3402/jev.v4.26316. https://doi.org/10.3402/jev.v4.26316.

[97] P. Rosa, F. Clementi, Absorption and Tissue Distribution of Doxorubicin Entrapped in Liposomes following Intravenous or Intraperitoneal Administration, PHA. 26 (1983) 221–229. https://doi.org/10.1159/000137805.

[98] G. Lee, S. Han, I. Inocencio, E. Cao, J. Hong, A.R.J. Phillips, J.A. Windsor, C.J.H. Porter, N.L. Trevaskis, Lymphatic Uptake of Liposomes after Intraperitoneal Administration Primarily Occurs via the Diaphragmatic Lymphatics and is Dependent on Liposome Surface Properties, Mol. Pharmaceutics. 16 (2019) 4987–4999. https://doi.org/10.1021/acs.molpharmaceut.9b00855.

[99] Z. Xueying, L. Zhelong, S. Wenqi, Y. Guodong, X. Changyang, Y. Lijun, Delivery Efficacy Differences of Intravenous and Intraperitoneal Injection of Exosomes: Perspectives from Tracking Dye Labeled and MiRNA Encapsulated Exosomes, Current Drug Delivery. 17 (2020) 186–194.

[100] H. Sun, D. Zhong, C. Wang, Y. Sun, J. Zhao, G. Li, MiR-298 Exacerbates Ischemia/Reperfusion Injury Following Ischemic Stroke by Targeting Act1, CPB. 48 (2018) 528–539. https://doi.org/10.1159/000491810.

[101] V. Boissonneault, I. Plante, S. Rivest, P. Provost, MicroRNA-298 and MicroRNA-328 Regulate Expression of Mouse β-Amyloid Precursor Protein-converting Enzyme 1 *, Journal of Biological Chemistry. 284 (2009) 1971–1981. https://doi.org/10.1074/jbc.M807530200.

[102] D. Barbagallo, S. Piro, A.G. Condorelli, L.G. Mascali, F. Urbano, N. Parrinello, A. Monello, L. Statello, M. Ragusa, A.M. Rabuazzo, C. Di Pietro, F. Purrello, M. Purrello, miR-296-3p, miR-298-5p and their downstream networks are causally involved in the higher resistance of mammalian pancreatic α cells to cytokine-induced apoptosis as compared to β cells, BMC Genomics. 14 (2013) 62. https://doi.org/10.1186/1471-2164-14-62.

[103] N. Bushati, S.M. Cohen, microRNA Functions, Annual Review of Cell and Developmental Biology. 23 (2007) 175–205. https://doi.org/10.1146/annurev.cellbio.23.090506.123406.

[104] X. Li, M. Commane, H. Nie, X. Hua, M. Chatterjee-Kishore, D. Wald, M. Haag, G.R. Stark, Act1, an NF-κB-activating protein, PNAS. 97 (2000) 10489–10493. https://doi.org/10.1073/pnas.160265197.

[105] V. Agarwal, G.W. Bell, J.-W. Nam, D.P. Bartel, Predicting effective microRNA target sites in mammalian mRNAs, ELife. 4 (2015) e05005. https://doi.org/10.7554/eLife.05005.

[106] Y. Chen, X. Wang, miRDB: an online database for prediction of functional microRNA targets, Nucleic Acids Research. 48 (2020) D127–D131. https://doi.org/10.1093/nar/gkz757.

[107] L.-L. Kong, X.-M. Zhuang, H.-Y. Yang, M. Yuan, L. Xu, H. Li, Inhibition of P-glycoprotein Gene Expression and Function Enhances Triptolide-induced Hepatotoxicity in Mice, Sci Rep. 5 (2015) 11747. https://doi.org/10.1038/srep11747.

[108] T. Kalpachidou, K.K. Kummer, M. Mitrić, M. Kress, Tissue Specific Reference Genes for MicroRNA Expression Analysis in a Mouse Model of Peripheral Nerve Injury, Front Mol Neurosci. 12 (2019) 283. https://doi.org/10.3389/fnmol.2019.00283.

[109] S. Mroczek, A. Dziembowski, U6 RNA biogenesis and disease association, WIREs RNA. 4 (2013) 581–592. https://doi.org/10.1002/wrna.1181.

[110] G. Lou, N. Ma, Y. Xu, L. Jiang, J. Yang, C. Wang, Y. Jiao, X. Gao, Differential distribution of U6 (RNU6-1) expression in human carcinoma tissues demonstrates the requirement for caution in the internal control gene selection for microRNA quantification, International Journal of Molecular Medicine. 36 (2015) 1400–1408. https://doi.org/10.3892/ijmm.2015.2338.

[111] N. Diette, J. Koo, S. Cabarcas-Petroski, L. Schramm, Gender Specific Differences in RNA Polymerase III Transcription, J Carcinog Mutagen. 7 (2016) 251. https://doi.org/10.4172/2157-2518.1000251.

[112] M.A. Pahlavani, M.D. Harris, The age-related changes in DNA binding activity of AP-1, NF-κB, and OCT-1 transcription factors in lymphocytes from rats, AGE. 19 (1996) 45–54. https://doi.org/10.1007/BF02434070.

[113] R. Ahuja, V. Kumar, Stimulation of Pol III-dependent 5S rRNA and U6 snRNA gene expression by AP-1 transcription factors, The FEBS Journal. 284 (2017) 2066–2077. https://doi.org/10.1111/febs.14104.

[114] C.D. Bryant, The blessings and curses of C57BL/6 substrains in mouse genetic studies, Ann N Y Acad Sci. 1245 (2011) 31–33. https://doi.org/10.1111/j.1749-6632.2011.06325.x.

[115] J.T. Eppig, J.A. Blake, C.J. Bult, J.A. Kadin, J.E. Richardson, The Mouse Genome Database (MGD): comprehensive resource for genetics and genomics of the laboratory mouse, Nucleic Acids Res. 40 (2012) D881–D886. https://doi.org/10.1093/nar/gkr974.

[116] R.E. Mebius, G. Kraal, Structure and function of the spleen, Nat Rev Immunol. 5 (2005) 606–616. https://doi.org/10.1038/nri1669.

[117] L.A. Medina, R. Klipper, W.T. Phillips, B. Goins, Pharmacokinetics and biodistribution of [111In]-avidin and [99mTc]-biotin-liposomes injected in the pleural space for the targeting of mediastinal nodes, Nuclear Medicine and Biology. 31 (2004) 41–51. https://doi.org/10.1016/S0969-8051(03)00122-7.

[118] K. Schwarz, M. Simons, J. Reiser, M.A. Saleem, C. Faul, W. Kriz, A.S. Shaw, L.B. Holzman, P. Mundel, Podocin, a raft-associated component of the glomerular slit diaphragm, interacts with CD2AP and nephrin, J Clin Invest. 108 (2001) 1621–1629. https://doi.org/10.1172/JCI12849.

[119] Polyclonal vs. monoclonal antibodies, (n.d.). https://www.ptglab.com/news/blog/polyclonal-vs-monoclonal-antibodies/ (accessed July 30, 2021).

[120] What is a polyclonal antibody? - Ximbio FAQ, (n.d.). https://ximbio.com/faq/57/what-is-a-polyclonal-antibody (accessed July 30, 2021).

[121] J.K. Hoober, ASGR1 and Its Enigmatic Relative, CLEC10A, Int J Mol Sci. 21 (2020) 4818. https://doi.org/10.3390/ijms21144818.

[122] J.D. Rosenblum, C.M. Boyle, L.B. Schwartz, The mesenteric circulation: anatomy and physiology, Surgical Clinics of North America. 77 (1997) 289–306. https://doi.org/10.1016/S0039-6109(05)70549-1.

[123] P. Towler, B. Staker, S.G. Prasad, S. Menon, J. Tang, T. Parsons, D. Ryan, M. Fisher, D. Williams, N.A. Dales, M.A. Patane, M.W. Pantoliano, ACE2 X-Ray Structures Reveal a Large Hinge-bending Motion Important for Inhibitor Binding and Catalysis*, Journal of Biological Chemistry. 279 (2004) 17996–18007. https://doi.org/10.1074/jbc.M311191200.

[124] A.J. Turner, Chapter 25 - ACE2 Cell Biology, Regulation, and Physiological Functions, in: T. Unger, U.M. Steckelings, R.A.S. dos Santos (Eds.), The Protective Arm of the Renin Angiotensin System (RAS), Academic Press, Boston, 2015: pp. 185–189. https://doi.org/10.1016/B978-0-12-801364-9.00025-0.

[125] M. Donoghue, F. Hsieh, E. Baronas, K. Godbout, M. Gosselin, N. Stagliano, M. Donovan, B. Woolf, K. Robison, R. Jeyaseelan, R.E. Breitbart, S. Acton, A novel angiotensin-converting enzyme-related carboxypeptidase (ACE2) converts angiotensin I to angiotensin 1-9, Circ Res. 87 (2000) E1–9. https://doi.org/10.1161/01.res.87.5.e1.

[126] D. Harmer, M. Gilbert, R. Borman, K.L. Clark, Quantitative mRNA expression profiling of ACE 2, a novel homologue of angiotensin converting enzyme, FEBS Letters. 532 (2002) 107–110. https://doi.org/10.1016/S0014-5793(02)03640-2.

[127] I. Hamming, W. Timens, M. Bulthuis, A. Lely, G. Navis, H. van Goor, Tissue distribution of ACE2 protein, the functional receptor for SARS coronavirus. A first step in understanding SARS pathogenesis, J Pathol. 203 (2004) 631–637. https://doi.org/10.1002/path.1570.

[128] F. Gembardt, A. Sterner-Kock, H. Imboden, M. Spalteholz, F. Reibitz, H.-P. Schultheiss, W.-E. Siems, T. Walther, Organ-specific distribution of ACE2 mRNA and correlating peptidase activity in rodents, Peptides. 26 (2005) 1270–1277. https://doi.org/10.1016/j.peptides.2005.01.009.

[129] S. Romagnani, T-cell subsets (Th1 versus Th2), Annals of Allergy, Asthma & Immunology. 85 (2000) 21. https://doi.org/10.1016/S1081-1206(10)62426-X.

[130] S. Kodidela, S. Ranjit, N. Sinha, C. McArthur, A. Kumar, S. Kumar, Cytokine profiling of exosomes derived from the plasma of HIV-infected alcohol drinkers and cigarette smokers, PLOS ONE. 13 (2018) e0201144. https://doi.org/10.1371/journal.pone.0201144.

[131] M.T. Abrams, M.L. Koser, J. Seitzer, S.C. Williams, M.A. DiPietro, W. Wang, A.W. Shaw, X. Mao, V. Jadhav, J.P. Davide, P.A. Burke, A.B. Sachs, S.M. Stirdivant, L. Sepp-Lorenzino, Evaluation of Efficacy, Biodistribution, and Inflammation for a Potent siRNA Nanoparticle: Effect of Dexamethasone Co-treatment, Molecular Therapy. 18 (2010) 171–180. https://doi.org/10.1038/mt.2009.208.

[132] F.A. Bonilla, H.C. Oettgen, Adaptive immunity, Journal of Allergy and Clinical Immunology. 125 (2010) S33–S40. https://doi.org/10.1016/j.jaci.2009.09.017.

[133] Q. Pan, V. Ramakrishnaiah, S. Henry, S. Fouraschen, P.E. de Ruiter, J. Kwekkeboom, H.W. Tilanus, H.L.A. Janssen, L.J.W. van der Laan, Hepatic cell-to-cell transmission of small silencing RNA can extend the therapeutic reach of RNA interference (RNAi), Gut. 61 (2012) 1330–1339. https://doi.org/10.1136/gutjnl-2011-300449.

[134] H. Nie, X. Xie, D. Zhang, Y. Zhou, B. Li, F. Li, F. Li, Y. Cheng, H. Mei, H. Meng, L. Jia, Use of lung-specific exosomes for miRNA-126 delivery in non-small cell lung cancer, Nanoscale. 12 (2020) 877–887. https://doi.org/10.1039/C9NR09011H.

[135] C. Liu, C. Su, Design strategies and application progress of therapeutic exosomes, Theranostics. 9 (2019) 1015–1028. https://doi.org/10.7150/thno.30853.

[136] G. Liang, Y. Zhu, D.J. Ali, T. Tian, H. Xu, K. Si, B. Sun, B. Chen, Z. Xiao, Engineered exosomes for targeted co-delivery of miR-21 inhibitor and chemotherapeutics to reverse drug resistance in colon cancer, Journal of Nanobiotechnology. 18 (2020) 10. https://doi.org/10.1186/s12951-019-0563-2.

[137] C. Meyer, J. Losacco, Z. Stickney, L. Li, G. Marriott, B. Lu, Pseudotyping exosomes for enhanced protein delivery in mammalian cells, IJN. 12 (2017) 3153–3170. https://doi.org/10.2147/IJN.S133430.

[138] S. Hsu, B. Yu, X. Wang, Y. Lu, C.R. Schmidt, R.J. Lee, L.J. Lee, S.T. Jacob, K. Ghoshal, Cationic lipid nanoparticles for therapeutic delivery of siRNA and miRNA to murine liver tumor, Nanomedicine: Nanotechnology, Biology and Medicine. 9 (2013) 1169–1180. https://doi.org/10.1016/j.nano.2013.05.007.

[139] Y. Akao, Y. Nakagawa, I. Hirata, A. Iio, T. Itoh, K. Kojima, R. Nakashima, Y. Kitade, T. Naoe, Role of anti-oncomirs miR-143 and -145 in human colorectal tumors, Cancer Gene Ther. 17 (2010) 398–408. https://doi.org/10.1038/cgt.2009.88.

[140] Y. Wu, M. Crawford, B. Yu, Y. Mao, S.P. Nana-Sinkam, L.J. Lee, MicroRNA Delivery by Cationic Lipoplexes for Lung Cancer Therapy, Mol. Pharmaceutics. 8 (2011) 1381–1389. https://doi.org/10.1021/mp2002076.

[141] Y. Wu, M. Crawford, Y. Mao, R.J. Lee, I.C. Davis, T.S. Elton, L.J. Lee, S.P. Nana-Sinkam, Therapeutic Delivery of MicroRNA-29b by Cationic Lipoplexes for Lung Cancer, Molecular Therapy - Nucleic Acids. 2 (2013) e84. https://doi.org/10.1038/mtna.2013.14.

[142] L. Jiang, H. Wang, S. Chen, Aptamer (AS1411)-Conjugated Liposome for Enhanced Therapeutic Efficacy of miRNA-29b in Ovarian Cancer, Journal of Nanoscience and Nanotechnology. 20 (2020) 2025–2031.

[143] Y. Chen, X. Zhu, X. Zhang, B. Liu, L. Huang, Nanoparticles Modified With Tumor-targeting scFv Deliver siRNA and miRNA for Cancer Therapy, Molecular Therapy. 18 (2010) 1650–1656. https://doi.org/10.1038/mt.2010.136.

[144] L. Wang, T.-T. Liang, CD59 receptor targeted delivery of miRNA-1284 and cisplatin-loaded liposomes for effective therapeutic efficacy against cervical cancer cells, AMB Express. 10 (2020) 1–11.

[145] R.W. Baker, B.T. Low, Membrane Separation⋆, in: Reference Module in Chemistry, Molecular Sciences and Chemical Engineering, Elsevier, 2015. https://doi.org/10.1016/B978-0-12-409547-2.11674-9.

[146] M. Sack, J. Sigler, S. Frenzel, Chr. Eing, J. Arnold, Th. Michelberger, W. Frey, F. Attmann, L. Stukenbrock, G. Müller, Research on Industrial-Scale Electroporation Devices Fostering the Extraction of Substances from Biological Tissue, Food Eng. Rev. 2 (2010) 147–156. https://doi.org/10.1007/s12393-010-9017-1.

[147] S. Kosuri, G.M. Church, Large-scale de novo DNA synthesis: technologies and applications, Nature Methods. 11 (2014) 499–507. https://doi.org/10.1038/nmeth.2918.

[148] J.F. Buyel, R.M. Twyman, R. Fischer, Very-Large-Scale Production of Monoclonal Antibodies in Plants, in: Process Scale Purification of Antibodies, John Wiley & Sons, Ltd, 2017: pp. 655–672. https://doi.org/10.1002/9781119126942.ch30.

[149] G. Liu, S. Hou, P. Tong, J. Li, Liposomes: Preparation, Characteristics, and Application Strategies in Analytical Chemistry, Crit Rev Anal Chem. (2020) 1–21. https://doi.org/10.1080/10408347.2020.1805293.

[150] B. Yu, R.J. Lee, L.J. Lee, Chapter 7 - Microfluidic Methods for Production of Liposomes, in: Methods in Enzymology, Academic Press, 2009: pp. 129–141. https://doi.org/10.1016/S0076-6879(09)65007-2.

[151] C. Charoenviriyakul, Y. Takahashi, M. Nishikawa, Y. Takakura, Preservation of exosomes at room temperature using lyophilization, International Journal of Pharmaceutics. 553 (2018) 1–7. https://doi.org/10.1016/j.ijpharm.2018.10.032.

[152] P.J. Stevens, R.J. Lee, Formulation kit for liposomal doxorubicin composed of lyophilized liposomes, Anticancer Res. 23 (2003) 439–442.

[153] Full article: Preparation and Characterization of Lyophilized Liposomes with Incorporated Quercetin, (n.d.). https://www.tandfonline.com/doi/full/10.1080/08982100500528594?casa_token=1GdStm2RxlIAAAAA%3AVRdurySrx65ftzPPDjrEC5mVUU6lETpUTD7hKIYpfJkuuunElVUblXKlm8mxS-fJG__pvQMo3RDF2g (accessed September 14, 2021).

[154] M. Glavas-Dodov, E. Fredro-Kumbaradzi, K. Goracinova, M. Simonoska, S. Calis, S. Trajkovic-Jolevska, A.A. Hincal, The effects of lyophilization on the stability of liposomes containing 5-FU, International Journal of Pharmaceutics. 291 (2005) 79–86. https://doi.org/10.1016/j.ijpharm.2004.07.045.

[155] T. Hernández-Caselles, J. Villalaín, J.C. Gómez-Fernández, Stability of Liposomes on Long Term Storage, Journal of Pharmacy and Pharmacology. 42 (1990) 397–400. https://doi.org/10.1111/j.2042-7158.1990.tb06578.x.

[156] P.R. Matias-Garcia, R. Wilson, V. Mussack, E. Reischl, M. Waldenberger, C. Gieger, G. Anton, A. Peters, A. Kuehn-Steven, Impact of long-term storage and freeze-thawing on eight circulating microRNAs in plasma samples, PLOS ONE. 15 (2020) e0227648. https://doi.org/10.1371/journal.pone.0227648.

[157] E.W. R., W. H., N. Y., C. J., Improving the Stability of Aptamers by Chemical Modification, Current Medicinal Chemistry. 18 (2011) 4126–4138.

[158] R. Crescitelli, C. Lässer, J. Lötvall, Isolation and characterization of extracellular vesicle subpopulations from tissues, Nat Protoc. 16 (2021) 1548–1580. https://doi.org/10.1038/s41596-020-00466-1.

[159] T.A. Hartjes, S. Mytnyk, G.W. Jenster, V. van Steijn, M.E. van Royen, Extracellular Vesicle Quantification and Characterization: Common Methods and Emerging Approaches, Bioengineering (Basel). 6 (2019) 7. https://doi.org/10.3390/bioengineering6010007.

[160] J.-H. Park, E.-W. Cho, S.Y. Shin, Y.-J. Lee, K.L. Kim, Detection of the Asialoglycoprotein Receptor on Cell Lines of Extrahepatic Origin, Biochemical and Biophysical Research Communications. 244 (1998) 304–311. https://doi.org/10.1006/bbrc.1998.8256.

[161] M.A. Saleem, M.J. O’Hare, J. Reiser, R.J. Coward, C.D. Inward, T. Farren, C.Y. Xing, L. Ni, P.W. Mathieson, P. Mundel, A Conditionally Immortalized Human Podocyte Cell Line Demonstrating Nephrin and Podocin Expression, JASN. 13 (2002) 630–638. https://doi.org/10.1681/ASN.V133630.

[162] D. Susan-Resiga, E. Girard, R. Essalmani, A. Roubtsova, J. Marcinkiewicz, R.M. Derbali, A. Evagelidis, J.H. Byun, P.F. Lebeau, R.C. Austin, N.G. Seidah, Asialoglycoprotein receptor 1 is a novel PCSK9-independent ligand of liver LDLR cleaved by furin, J Biol Chem. 297 (2021) 101177. https://doi.org/10.1016/j.jbc.2021.101177.

[163] C.G. Gaetano, N. Samadi, J.L. Tomsig, T.L. Macdonald, K.R. Lynch, D.N. Brindley, Inhibition of autotaxin production or activity blocks lysophosphatidylcholine-induced migration of human breast cancer and melanoma cells, Mol Carcinog. 48 (2009) 801–809. https://doi.org/10.1002/mc.20524.

[164] S.Y. Yang, J. Lee, C.G. Park, S. Kim, S. Hong, H.C. Chung, S.K. Min, J.W. Han, H.W. Lee, H.Y. Lee, Expression of autotaxin (NPP-2) is closely linked to invasiveness of breast cancer cells, Clin Exp Metastasis. 19 (2002) 603–608. https://doi.org/10.1023/A:1020950420196.

[165] R. Leblanc, S.-C. Lee, M. David, J.-C. Bordet, D.D. Norman, R. Patil, D. Miller, D. Sahay, J. Ribeiro, P. Clézardin, G.J. Tigyi, O. Peyruchaud, Interaction of platelet-derived autotaxin with tumor integrin αVβ3 controls metastasis of breast cancer cells to bone, Blood. 124 (2014) 3141–3150. https://doi.org/10.1182/blood-2014-04-568683.

[166] B.-J. Zhai, Z.-Y. Shao, C.-L. Zhao, K. Hu, F. Wu, Development and characterization of multidrug resistant human hepatocarcinoma cell line in nude mice, World J Gastroenterol. 12 (2006) 6614–6619. https://doi.org/10.3748/wjg.v12.i41.6614.

[167] M.W. Pfaffl, A. Tichopad, C. Prgomet, T.P. Neuvians, Determination of stable housekeeping genes, differentially regulated target genes and sample integrity: BestKeeper – Excel-based tool using pair-wise correlations, Biotechnology Letters. 26 (2004) 509–515. https://doi.org/10.1023/B:BILE.0000019559.84305.47.

[168] N. Silver, S. Best, J. Jiang, S.L. Thein, Selection of housekeeping genes for gene expression studies in human reticulocytes using real-time PCR, BMC Mol Biol. 7 (2006) 33. https://doi.org/10.1186/1471-2199-7-33.

[169] J. Vandesompele, K. De Preter, F. Pattyn, B. Poppe, N. Van Roy, A. De Paepe, F. Speleman, Accurate normalization of real-time quantitative RT-PCR data by geometric averaging of multiple internal control genes, Genome Biol. 3 (2002) research0034.1-research0034.11.

[170] C.L. Andersen, J.L. Jensen, T.F. Ørntoft, Normalization of Real-Time Quantitative Reverse Transcription-PCR Data: A Model-Based Variance Estimation Approach to Identify Genes Suited for Normalization, Applied to Bladder and Colon Cancer Data Sets, Cancer Res. 64 (2004) 5245–5250. https://doi.org/10.1158/0008-5472.CAN-04-0496.

[171] ICMJE | Recommendations | Defining the Role of Authors and Contributors, (n.d.). http://www.icmje.org/recommendations/browse/roles-and-responsibilities/defining-the-role-of-authors-and-contributors.html (accessed July 29, 2020).

[172] D.B. Resnik, A.M. Tyle, J.R. Black, G. Kissling, Authorship policies of scientific journals, J Med Ethics. 42 (2016) 199–202. https://doi.org/10.1136/medethics-2015-103171.

[173] E.E. Tarkang, M. Kweku, F.B. Zotor, Publication Practices and Responsible Authorship: A Review Article, J Public Health Afr. 8 (2017). https://doi.org/10.4081/jphia.2017.723.

[174] K. Strange, Authorship: why not just toss a coin?, Am J Physiol Cell Physiol. 295 (2008) C567–C575. https://doi.org/10.1152/ajpcell.00208.2008.

